# cAMP binding to closed pacemaker ion channels is cooperative

**DOI:** 10.1101/2023.04.18.537331

**Authors:** Stefan Kuschke, Susanne Thon, Tina Schwabe, Klaus Benndorf, Ralf Schmauder

## Abstract

Most of receptor proteins in biomembranes are oligomers of either homologue or identical subunits, that govern a cooperative activation process. HCN ion channels, generating electric rhythmicity in specialized brain neurons and cardiomyocytes^1^, are tetramers activated by hyperpolarizing voltage and modulated by the intracellular binding of the second messenger cAMP^2,3^. The molecular processes underlying the cooperative action of the subunits^4-6^ by the binding are not fully understood, in particular if they are necessarily associated with channel activation^7^ or not, as recently reported for detergent-solubilized receptors positioned in zero-mode waveguides^8^. Here we show positive cooperativity in ligand binding to resting HCN2 channels in native cell membranes by analyzing the binding of fluorescence-labelled single cAMP derivatives to the channels. Kinetic modelling reveals, that the affinity of the still empty binding sites rises with increased degree of occupation and that the transition of the channel to a flip state is promoted accordingly. We conclude that ligand binding to the subunits in resting HCN2 channels is already cooperative prior to channel activation. This shows that single-molecule binding studies can quantify cooperativity in ligand binding to receptors in native membranes based on equilibrium measurements.

---

Hyperpolarization-activated cyclic nucleotide-modulated (HCN-)channels are nonspecific cation channels and they belong to the superfamily of six-transmembrane domain voltage-gated channels^9-11^. The obligatory activation stimulus of these channels is a sufficiently negative membrane voltage. In addition, the binding of the second messenger cAMP to the four available intracellular cyclic nucleotide-binding domains (CNBDs), one in each subunit, boosters activation by both shifting activation to less negative voltages and accelerating the activation speed^6^.

The concentration-activation relationship of HCN2 channels with cAMP can be approximated by a Hill function with Hill coefficients exceeding unity^12^, indicating a cooperative effect of cAMP binding on activation. Combined current and binding measurements in ensembles of channels by confocal patch-clamp fluorometry (cPCF)^13^ suggested cooperativity with the sequence positive-negative-positive^4,5^. This cooperativity might emerge from a combination of events from ligand binding to channel activation or from activation only by a reciprocal feedback from activation to ligand binding^12^.

Results from cPCF measurements on ligand binding to resting HCN2 channels in patches with ensembles of channels required Hill coefficients larger than unity^14,15^, ruling out independent and supporting cooperative binding to the subunits. In contrast, a recent single-molecule binding study reported that cAMP binds independently to all four subunits in both closed HCN1 and HCN2 channels^8^. The authors performed single-molecule binding measurements with a fluorescent cAMP derivative to detergent solubilized receptors positioned in zero-mode waveguides (ZMWs). These experiments suggested also the existence of a flip state following the binding^15^. Together, these data favor a scenario that only hyperpolarization-induced channel activation but not cAMP binding alone evokes relevant cooperative interaction of the subunits, leaving the elevated Hill coefficient in macroscopic binding experiments^14,15^ unexplained.

Single molecule detection has become a powerful technique to elucidate details of molecular processes^16,17^ at equilibrium. However, most of single-molecule experiments do not directly follow ligand-receptor interaction though it is the initiating step of most signaling cascades^18^. In some studies ligands were used for continuous reversible labelling in single-molecule tracking^18,19^ and super-resolution microscopy^20^. However, single-molecule binding studies are rare, because only few suitable fluorescent agonists are known and the signal-to-noise ratio is limited due to signals from unbound ligands. Several strategies were employed to increase the signal-to-noise ratio: (1) Selection of high affinity systems by mutations or using either natural^21,22^ or engineered^23,24^ ligands with higher affinities. (2) Reduction of the detection volume either mechanically^25^ or optically, e.g. with total internal reflection Microscopy (TIRFM)^26^ or confocal techniques as fluorescence correlation spectroscopy (FCS)^27,28^. A particular case are ZMWs^29^ that were successfully used to investigate ligand binding on isolated binding sites^30,31^ or solubilized channels^8^. So far, there are only proof-of-concept experiments with ligand binding to native membranes in planar waveguides^32^. (3) Utilization of brighter fluorophores to reduce relative contributions of the Poisson noise^33^.

Here we use a combination of these strategies to enable single-molecule binding measurements: We use a ligand with both increased affinity and brightness compared to previous studies^8,15^ and employed TIRFM to reduce signals from ligands in solution. We follow the binding of single ligands to individual mHCN2 channels embedded in a native membrane. We demonstrate positive cooperativity in ligand binding to these channels contrasting a recent study in solubilized receptors^8^. The consecutive binding steps increase both the ligand affinity of the binding sites and the population of a flip state.

## f1cAMP binding to HCN2 channels

We established a stable HEK 293 cell line expressing N-terminally GFP-tagged HCN2 channels (eGFPmHCN2) using an inducible promoter. The eGFP-HCN2 construct showed voltage activation and ligand modulation comparable to wildtype HCN2 (Extended Data Fig. 1). For visualization of ligand binding, a protocol similar to that of Perez and coworkers^34^ was established, yielding native membrane sheets with the intracellular side exposed to solution.

Specific binding of f1cAMP to the GFP-tagged HCN2 channels was confirmed (Extended Data Fig. 4c). The used f1cAMP allowed for reliable single-molecule detection at concentrations up to 0.5 μM in TIRFM. Thus, at controlled low expressions (0.2 channels/μm^2^), single-ligand binding dynamics to individual HCN2 channels could be followed.

While the environment of a native membrane excludes artefacts from channel isolation and solubilization, its complexity can distort the measurements by ligands binding to endogenous binding sites. We reduced the potential background by competitive suppression^35-37^ of spurious nonspecific binding to endogenous cAMP binding proteins^38-40^ (Extended Data Fig. 5) and ruled out diffusional dynamics on the time and length scale of our analysis (Extended Data Fig. 2). This ensured that our experiments report only binding to eGFPmHCN2 channels and not binding to endogenous proteins or diffusional fluctuations.

Intensity trajectories for each identified mHCN2 channel were extracted, filtered, and idealized (Fig. 2a). As a model-free approach to detect cooperativity, observed occupancies were compared to the binomial expectations for four independent binding sites (Fig. 2c). For concentrations of 0.3 and 0.5 μM f1cAMP, only the double liganded state matched the expected fractions. The non-liganded and the highly liganded states with 3 and 4 ligands bound were more abundant where the first liganded state was less abundant than the binomial expectations, contradicting independent binding^8^. Notably, more complex models of independent binding sites would not alter these equilibrium binding probabilities, and only these and not dynamics are considered in this analysis.

**Fig. 1.**
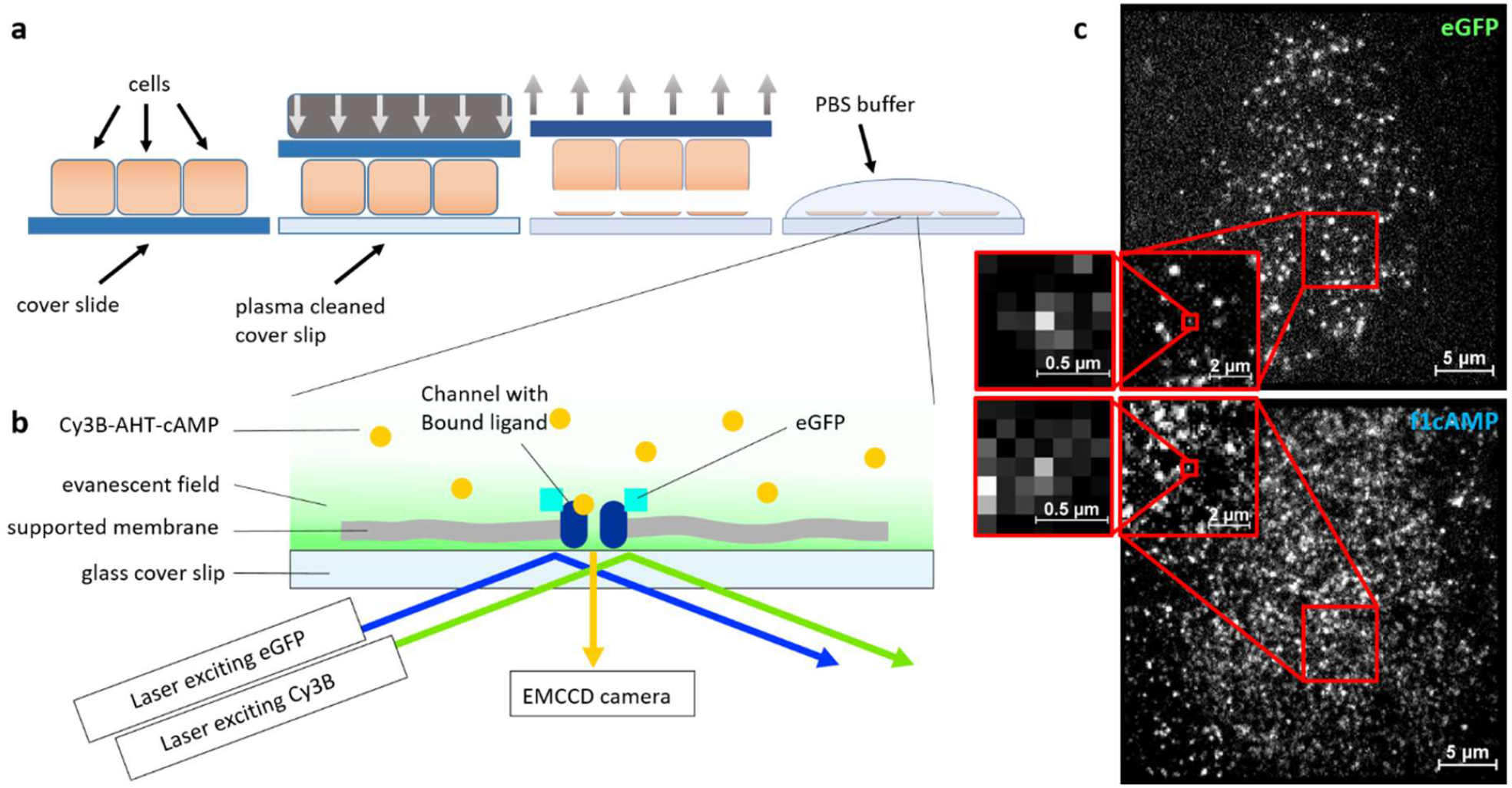
Generation of supported membrane sheets. **a**, Supported membrane sheets were generated by using cultured HEK293 cells stably transfected with eGFPmHCN2 and sticking the membrane to cover slips. Cover slips were put upside down onto plasma-cleaned cover slides and pushed down by 5.3 kPa. Cover slips were then removed, leaving the top membrane on the cover slide while the intracellular part was accessible for solutions. **b**, TIRF measurements of fluorescent cAMP derivatives and eGFP tags attached to HCN2 channels were performed on supported membranes. Only ligands near the membrane are excited due to the typical steep exponential decay of the laser power in the evanescent field. **c**, eGFP (top) and f1cAMP (bottom) TIRF recordings of a supported membrane. Individual signals were fitted by a 2D Gaussian function to ensure evaluation of single molecules.

**Fig. 2.**
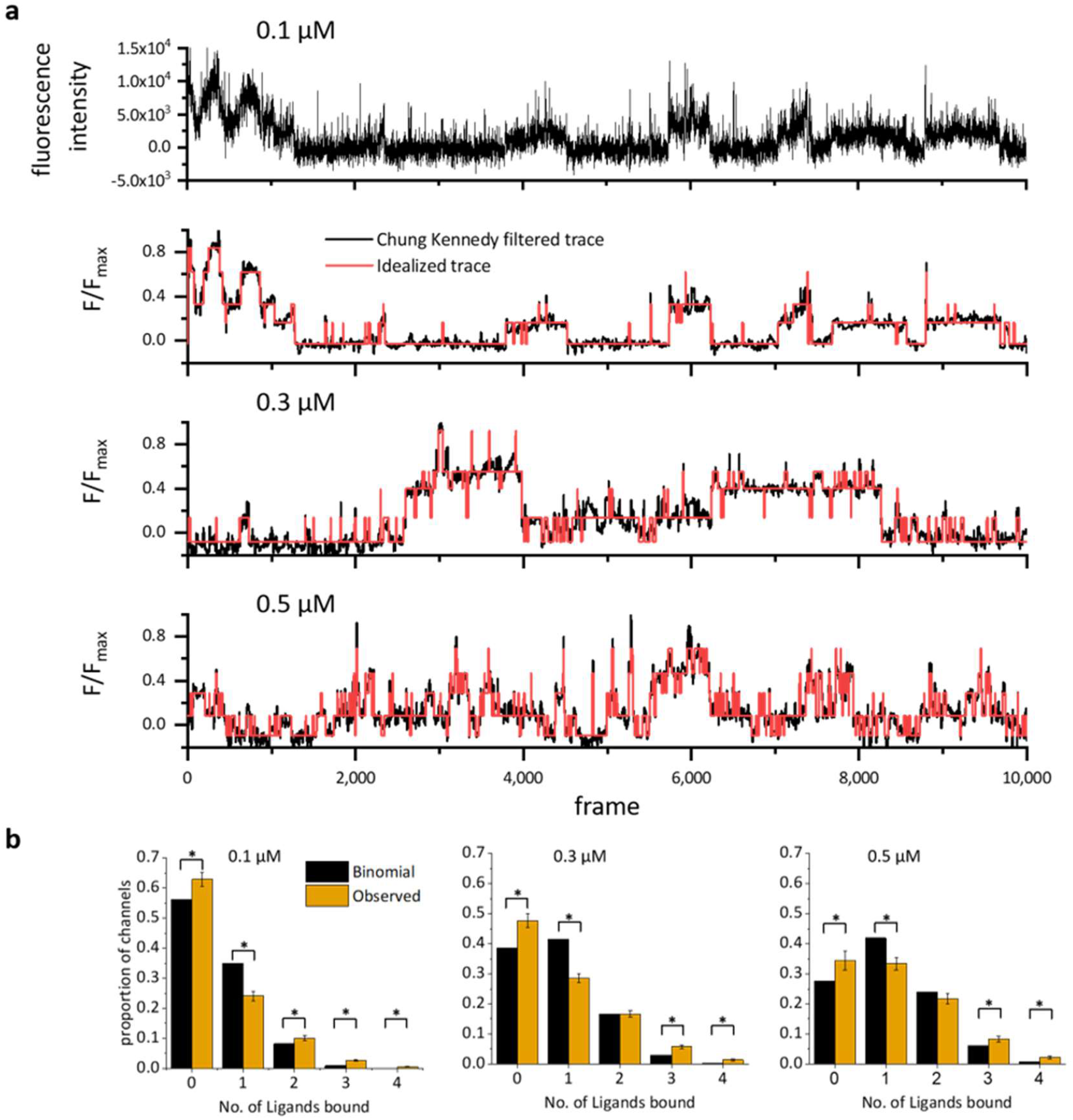
Recording of binding events and data processing. **a**, Raw data traces were smoothened by the Chung-Kennedy Filter. Filtered traces were idealized to the number of bound ligands by the DISC algorithm^35^. **b**, The proportion of observed states was compared to the expected proportion of non-cooperative binding calculated by equation (2). Error bars show sem. *indicate significant differences (one sample *t*-test, P<0.05).

To further interpret our results, we first determined the transition rates for a simple model that includes four consecutive binding steps and stoichiometric factors for binding and unbinding (Fig. 3a, Extended Data Table 2, model 12). Binding and unbinding rate constants (Fig. 3b,c) were higher than previously reported, likely due to the higher affinity of f1cAMP compared to the previously used fcAMP^23^. The association constants (Fig. 3d) showed an almost exponential increase along the 4 binding steps. This differs from the data shown by White and coworkers^8^ where association constants remain almost constant. It also differs from data by Thon and coworkers^15^ that showed an initial decrease after the first binding step, followed by a strong increase for the last binding step. However, these models differed in structure and assumptions.

**Fig.3.**
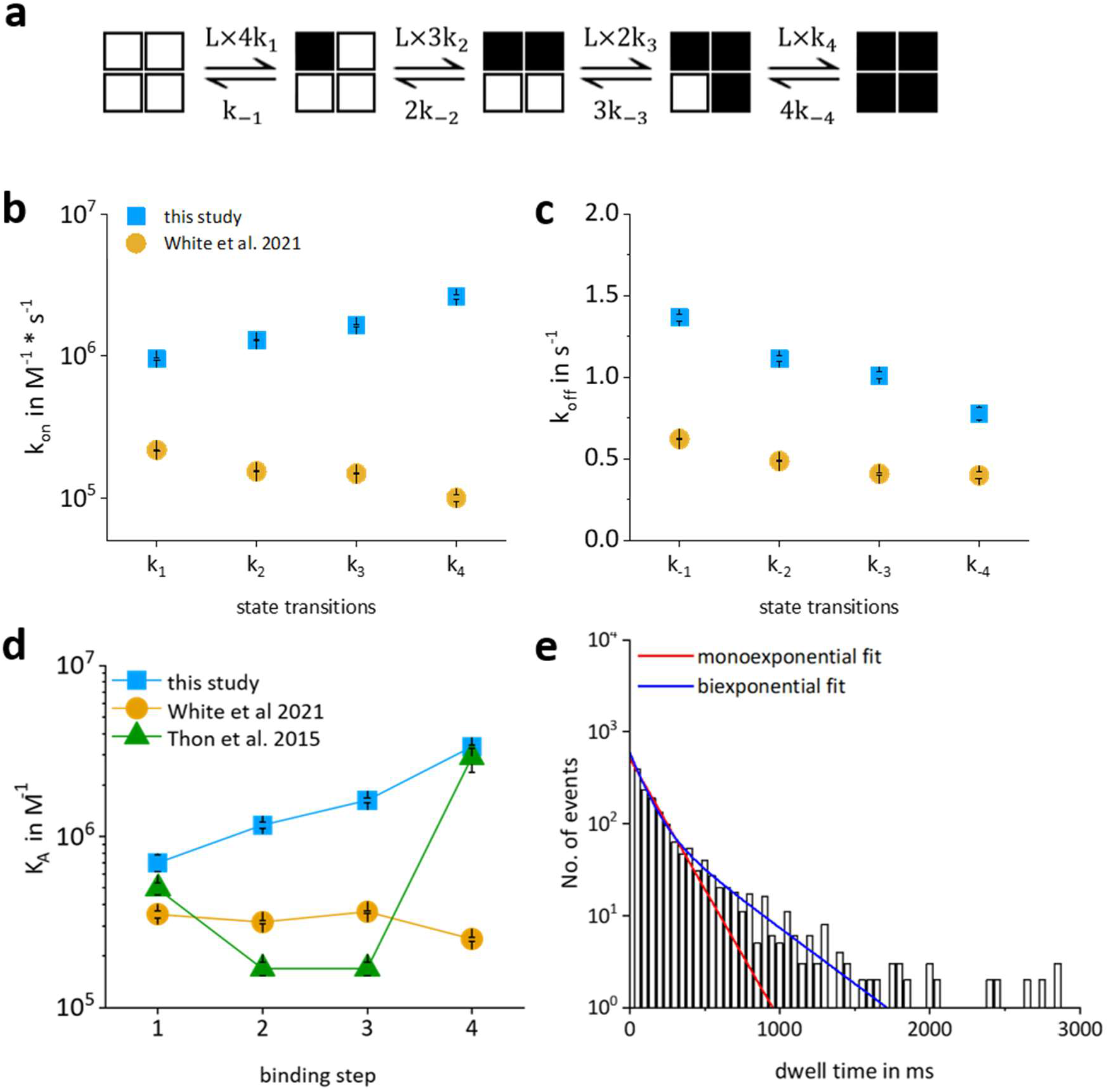
Kinetic and equilibrium constants calculated for a simple binding reaction. **a**, simplest model with four consecutive binding steps **b-d**, Comparison of present to previously published constants. **b**, microscopic binding rate constants *k*_*on*_ and **c**, microscopic dissociation rate constants *k*_*on*_ as well as **c**, association constants *K*_*A*_ of individual binding steps. **d**, Dwell times of isolated single-liganded states at 0.1 μM f1cAMP fitted by a mono- or biexponential function.

### Ligand binding populates a flip state

Dwell times of selected single-liganded states that were both preceded and followed by non-liganded states were extracted. For all concentrations, the necessity of a second time constant (Fig. 3e and Extended Data Fig.2), indicates directly that ligand binding promotes occupation of another state, termed in the following flip state.

To understand the dynamics, we analyzed in total 13 kinetic models (Extended Data Materials, Extended Data Table 2). Taking both the tetrameric structure with four binding sites and the existence of a flip state into account, we started the modeling with schemes containing 10 states, five resting states and five flip states in a parallel arrangement (model 1 in Extended Data Table 2). We applied the principle of microscopic reversibility, reducing the number of rate constants per cycle by one. The log-likelihood (LL) value was taken as measure for the goodness of the fit, the parameter uncertainty (relative SEM) was considered as well (Extended Data Table 3).

Following the strategy described in Extended Data Methods, model 12 combined the highest likelihood and lowest uncertainty (Fig. 4a, Extended Data Table 2). From the rate constants of the individual steps, equilibrium association constants, K_A_=k_x_/k_-x_, and equilibrium flip constants, K_F_=f_x_/f_-x_, were calculated. For the resting states, K_A_, increases in a non-linear fashion from the first to the fourth binding step (Fig. 4b). In the flip state, the K_A_ values are larger for the binding steps 2, 3, and 4 with respect to their counterparts in the resting state by about threefold. Similar to the resting state, the association rate constant stays almost constant for binding step 3 with respect to binding step 2 but almost doubles for binding step 4 with respect to binding step 3. Regarding the flip reaction, the empty and the single-liganded state have about the same K_F_ value whereas the binding of the ligands 2, 3, and 4 increases K_F_ strongly in an approximately exponential fashion (Fig. 4c). These results show that in channels not pre-activated by voltage, the first binding step is without major effect. In contrast, the second binding step plays a key role for triggering an enhanced ligand binding in the third and fourth binding step.

**Fig. 4.**
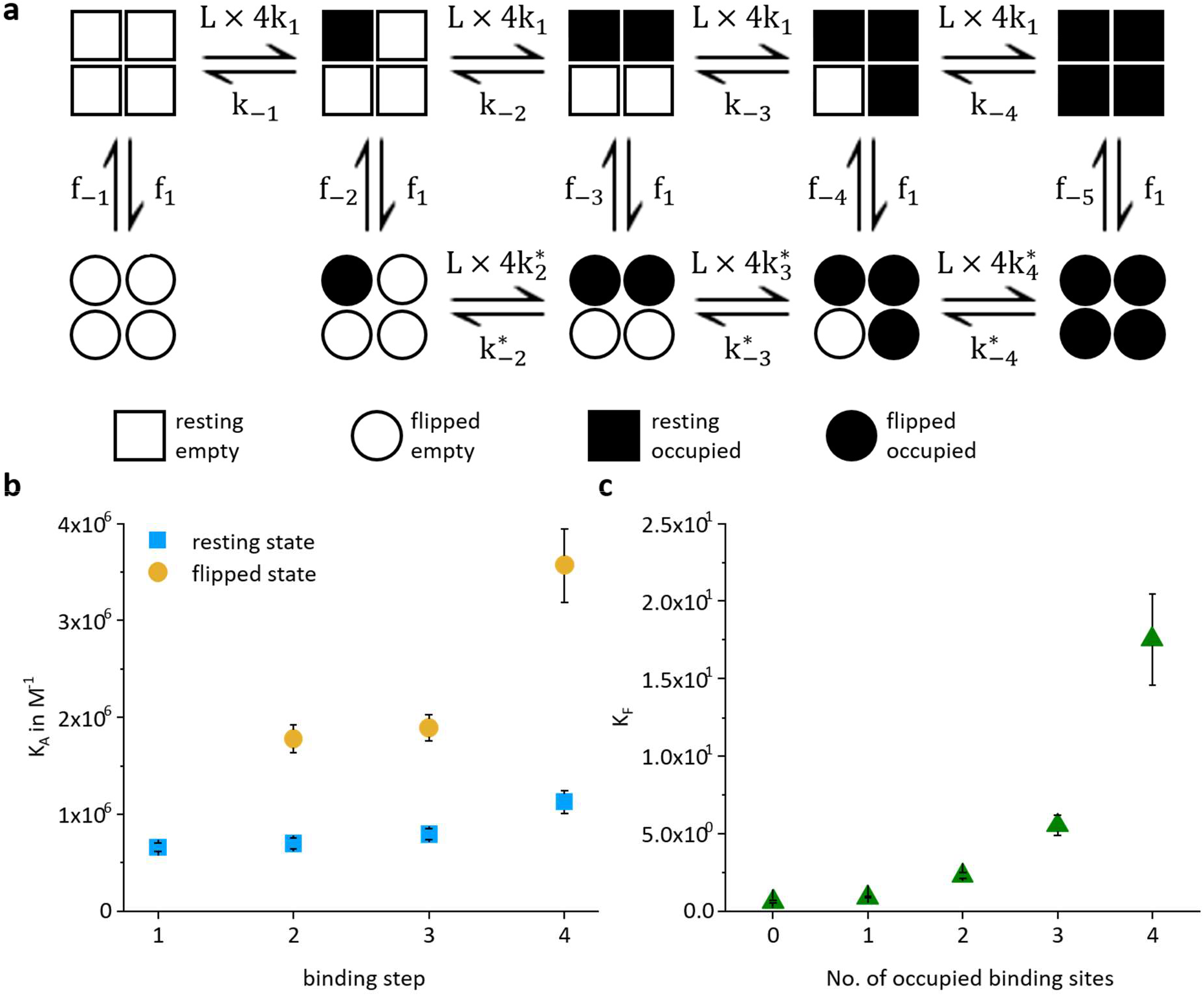
Favored model to describe the single binding events. **a**, Scheme of model 13. The model consists of four individual binding steps of which each has an own transition to a flip state. In a flip state, additional binding is possible. Due to extremely small rate constants between the non-liganded and the single-liganded flip state, this transition is excluded. Symbols describe subunit properties. **b**, Equilibrium association constants (K_A_) for binding of ligands in the non-flip (blue) and flip state (orange). **c**, Equilibrium constants for switching into the flip state depending on the state of receptor occupation. Error bars show sem.

## Discussion

Here we quantified the kinetics of ligand-binding on single resting tetrameric mHCN2 channels, functionally expressed in native cell membranes. We demonstrate that the binding process in the resting channels is cooperative and associated with a molecular flip reaction. These results are in contrast to a previous study on single-molecule binding by White and coworkers^8^ but are in line with results on previous macroscopic measurements on the kinetics of ligand binding in excised patches by Thon and coworkers^15^.

To explain these differences, we compare data collection and analysis in more detail: White and coworkers established the DISC algorithm to analyze data^35^, which we adapted to our conditions (Extended Data Adaptation of DISC – Algorithm). In both single-molecule studies, eGFP-tagged HCN2 channels were used whereas untagged channels were used by Thon and coworkers^15^. In previous studies the fluorescent cAMP derivative fcAMP (Dy547-AET-cAMP) was used while we used the superior f1cAMP (Cy3B-AHT-cAMP)^23^ that shows an about 2.5 times higher affinity for HCN2 channels than fcAMP^14,23^ and contains a brighter fluorophore, allowing for faster data acquisition. This allows to use lower ligand concentrations, enabling to employ conventional TIRFM, while White and coworkers recorded in zero-mode wave guides.

Another major difference lies in the sample preparation and, following this, the molecular environment of the channels: White and coworkers^8^ solubilized and purified the channels, into artificial detergent micelles whereas we studied the binding in native cell membranes. While micelles contain only one protein, the mHCN2 channel, our native membrane contains also other proteins potentially binding f1cAMP. To minimize inclusion of binding to these, we employed two strategies: (I) We utilized eGFP tagged channels and only evaluated binding signals with point-spread-function (PSF) like shapes that where closely colocalized to the PSF-shaped eGFP signal. (II) We reduced the influence of endogenous binding sites^36-38^ by adding a mixture of selective competitive inhibitors (Extended Data Methods)^39-41^. The functionality of the mHCN2 channels was not altered by these inhibitors (Extended Data Fig. 4). Taken together, we suggest that the functional difference between our results and those of White and coworkers – cooperativity versus non-cooperativity in ligand binding – arise from the different nature of the membranes surrounding the channel.

Relevant influences of the membrane composition of micelles on either the structure or function of membrane proteins is a well-known phenomenon. For example, Cao and coworkers^42^ showed for mGlu2 receptors that the interaction between membrane forming lipids and the receptor is required for function. With regard to HCN channels, a direct context between membrane composition and channel function was reported by Fürst and D’Avanzo^43^ who investigated the role of cholesterol on activation kinetics, deactivation kinetics and current density in cells. Though Fürst and D’Avanzo^43^ did not consider cooperativity phenomena, their results with cholesterol enriched membranes might result from cooperativity changes. Interestingly, White and coworkers^8^ had to optimize the detergent condition to obtain stable sample preparations for different HCN-isoforms, which always included a steroidal detergent.

To quantify the mode of cooperativity we fitted a series of kinetic models and subsequently adopted the following three assumptions: (1) Binding rates to resting channel subunits were set to be equal after accounting for stoichiometry. (2) Rates of entering the flip state were linked to increase the accuracy of the determined parameters while off-rates were kept independent to account for possible ligand-protein interactions stabilizing the flip state. (3) Binding to neighboring subunits was not distinguished from that to opposite subunits and among the neighboring subunits also not between the right and the left neighbor. We assume this is justified as no respective difference was reported for channel activation in mutated concatamers^44^. With these assumptions the best model was chosen based on the certainty of the estimated parameters and the LL. Binding rates were comparable to values published for related isolated binding sites^45^.

Together, our modelling for resting mHCN2-channels suggests (I) cooperativity in ligand binding, (II) a transition to a flip state that is favored with increased occupation of the binding sites, and (III) that ligand binding is possible in both resting and flipped channels. In our considerations, the flip transition is assigned to the whole channel, as e.g. in the Monod-Wyman-Changeux model^46^.

While standing in contrast to data from White and coworkers^8^ our study agrees with data reported by Thon and coworkers^15^, concerning the existence of cooperativity, however the observed cooperativity differs. A flip state was hypothesized by both groups^15,8^. We investigated equilibrium fluctuations of f1cAMP-binding in HEK293 membranes, while Thon and coworkers^15^ followed binding upon fcAMP concentration jumps on *Xenopus* oocyte patches.

In conclusion, our data demonstrate that the subunits in resting HCN2 channels act in a cooperative fashion when ligands bind to the four binding sites and that this cooperative action of the subunits is associated with the transition to a flip state. In general this indicates that cooperativity emerges already upon ligand-binding and does not require activation of the channel pore.

## Methods

### Single molecule binding measurements

Briefly, HEK293 derived cells expressing eGFPmHCN2 were grown on Poly-L-lysine coated cover-slips, inverted onto plasma clean cover-slides, pressed on, and removed, yielding supported native membrane patches on the cleaned coverslides. These were mounted on an Eclipse Ti inverted Microscope and imaged in TIRF (1.49/100x) onto an Andor 888 Ultra EMCCD in isolated crop mode. Cells were incubated with f1cAMP supplemented with an inhibitor mix^39-41^ and an oxygen scavenger system. Subsequent movies of ligand binding dynamics were recorded with 542 nm excitation followed by a movie with 488 nm excitation identifying channel positions by single-step bleaching of the eGFPs.

### Single molecule analysis

Time trajectories of single molecules binding and unbinding were extracted and processed using MATLAB 2020b (Mathworks). Background and drift were removed by subtracting a median filtered movie (Kernel: 9 × 9 × 3). Positions of binding and unbinding events were matched to positions of eGFP bleaching signals to select the positions of mHCN2 channels. 3×3 pixel ROIs were summed, generating time-traces corresponding to binding signal at individual mHCN2-channel (see: Extended methods) Traces of f1cAMP and eGFP signals were included if they showed stepwise changes in average intensity. Traces were Chung-Kennedy filtered^47,48^ and idealized using a modified version of the DISC algorithm^35^ (Extended Data Adaptation of DISC – Algorithm). All idealized traces were inspected by eye following idealization. Accepted traces accumulated to 1.6 × 10^4^ s of measurements across approximately 150 mHCN2 channels at f1cAMP concentrations of 0.1, 0.3 and 0.5 μM. From these traces population of liganded states, average binding probability and dwell times were extracted

### Modeling

Models (Extended Data Table 1) were optimized globally over all concentrations using maximum idealized point optimization^49^ as implemented in QuB^50^. The quality of optimization was evaluated by ranking the maximum likelihood and relative errors of the parameters.

## Supporting information

Suplementary data, methodes and discussion

## Author contributions

K.B. and R.S. envisioned the project, R.S. and S.T. established the single molecule measurement technique, S.K. performed the single molecule measurements, T.S. engineered the HEK293 cell line with inducible eGFPmHCN2 channel, S.K. and S.T. performed the patch clamp measurements, S.K. and R.S. wrote custom scripts and analyzed data, K.B., S.K., and R.S. wrote the manuscript

## Acknowledgement

We thank K. Schoknecht, S. Bernhardt and C. Ranke for technical assistance. We are grateful to A. Schweinitz, U. Enke and M. Lelle for the synthesis of the fluorescent ligand. The work was funded by the DFG (Transregio 166, project A5 to K.B. and R.S.; RU DynIon 2518, project P2, to K.B.).

