## Supplementary material for "cAMP binding to closed pacemaker ion channels is cooperative": Suplementary data, methodes and discussion

### Supplementary Methods

#### Chemicals

Chemicals were purchased and used with no further purification: 8-pCPT-2'OMe-cAMP (Biolog LSI GmbH & Co KG, Bremen, Germany), Trolox, Piclamilast, Glucose oxidase, Catalase, Glucose, Poly-L-Lysin (Merck KGaA, Darmstadt, Germany), ht31 was synthesized on demand (Genosphere Biotechnologies, Clamart, France), Cy3B-AHT-cAMP was synthesized in house as described previously[1].

#### Cell Culture

HEK293 cell lines containing an inducible promoter (Flp-In-T-REx 293; Invitrogen #R78007) were cultured in minimum essential medium (MEM, (Thermo Fisher Scientific, Schwerte, Germany)) supplemented with 10% fetal calf serum (BioWest, Nuaillé, France), non-essential amino acids (Gibco) and antibiotics according to the manufacturer's instructions. For stable cell lines, Flp-In-T-REx 293 cells were transfected using the calcium phosphate method with a mixture of plasmids (0.5 µg eGFPmHCN2 and 1.5 µg pOG44). After hygromycin B treatment, stable clones were selected and cultured until a passage number of approximately 20. eGFPmHCN2 contains an eGFP linked by a 6-amino acids linker (SDPNST) to mHCN2 (NM_008226)[2].

For single molecule measurements, cells were plated on cover slips (13 mm diameter, #1 - 0.15 mm thickness, Menzel Gläser, Braunschweig, Germany) in 24 well plates. Cover slips were coated with poly-L-lysin before use (0.1 mg/mL poly-L-lysine solution for 2 h followed by multiple washes with media). Per cover slip, 200 µL of resuspended cells (500 cells/µL) culture were added. Cells were allowed to settle a day before induction of protein expression with media supplemented with 0.2 µg/mL tetracycline (Sigma-Aldrich, Taufkirchen, Germany). Cells were used 2 to 48 h after induction.

#### Membrane preparation

Cells grown on cover slips were washed three times with sterile filtered PBS (300 µL). Cover slides (25 mm diameter, #1 - 0.15 mm thickness, Menzel Gläser, Braunschweig, Germany), prior to usage cleaned by Zepto Plasma cleaner (Diener Electronics, Ebhausen, Germany), were mounted in Attofluor imaging chambers (Thermo Fisher Scientific, Schwerte, Germany). Cover slips were inverted onto plasma cleaned cover slides with 20 µL PBS in-between the glasses. Additional weight (0.7 N) was put onto the glass slips for 5 minutes. After taking off the weight, the upper glass slip was removed using an EDSYN LP200 Pixter Pen-Vac pneumatice forceps (EDSYN Inc., Van Nuys, USA) leaving supported membrane attached to the lower, plasma cleaned cover slide. The cover slide with the attached membrane was washed three additional times with sterile filtered PBS. A solution of 100 µM ht31 in sterile filtered PBS was added and replaced after 5 minutes with sterile filtered PBS. Afterwards, membranes were incubated with a solution of f1cAMP (Cy3B-AHT-cAMP)[1,3], competitive inhibitors for membrane associated cAMP binding proteins (8-pCPT-2'OMe-cAMP, Piclamilast), an oxygen scavenger system (165 U/mL Glucose oxidase, 2170 U/mL Catalase, 0.4 w% Glucose) and 2 mM Trolox. The system was allowed to equilibrate for 5 minutes before imaging.

**Staining of membranes with DiO**

Vybrant™ DiO stock solution (Invitrogen Life Technologies, Eugene, Oregon, USA) was heated in supersonic bath for 15 minutes prior to use. The solution was diluted to 5 µM and 330 µL were added to each well containing HEK293 cells in a 24 – well plate (Greiner Bio-One, Frickenhausen, Germany). Cells were incubated at 37 °C with 5 % $CO_2$ for 10 minutes. Cells were washed three times with 300 µL PBS. From these cells, membranes were prepared according to Methods: Membrane preparation and used to determine background binding of f1cAMP.

**Oocyte preparation**

Oocytes were collected from adult *Xenopus laevis* through operation under anesthesia (1.2 g Tricaine mesylate, 2.4 g bullrich salt in 1L water). Oocytes were treated with 1.5 mg/mL collagenase (Roche, Grenzach-Wyhlen, Germany) for 105 minutes and manually dissected and injected with mRNA encoding wild type mHCN2 or eGFPmHCN2. After injection, oocytes were kept at 18 °C for 1-5 days in Barth's medium. The surgery procedures were carried out in accordance with the German Animal Welfare Act with the approval of the Thuringian State Office for Consumer Protection on 30.08.2013 and 09.05.2018.

**Patch-clamp recordings**

Currents were recorded in inside-out patches. Patch pipettes were pulled from quartz tubing with resistances of about 1.5 MΩ. Bath solution contained 100 mM KCl, 10 mM EGTA, 10 mM HEPES buffered to pH=7.4. Pipette solution contained 120 mM KCl, 10 mM HEPES, 1 mM $CaCl_2$. For maximum channel open probability, the patch was put into a stream of bath solution containing 10 µM cAMP. Currents were recorded with an Axopatch 200B amplifier (Axon Instruments, Foster City, CA). Rundown was avoided by waiting for 5 minutes after obtaining a gigaseal. Stimulation and data recording were performed with the ISO3 hard- and software (MFK, Niedernhausen, Germany) and with PATCHMASTER and LIH 8+8 hard- and software (HEKA Elektronik Dr. Schulze GmbH, Lambrecht, Germany). The sampling rate was 5 kHz and the on-line filter (four-pole Bessel) was set to 1 kHz.

**Single Molecule Microscopy**

Membranes were positioned first in the widefield of a microscope and then investigated by TIRFM. Measurements were performed on a Nikon Eclipse Ti inverse microscope using an Apochromat 1.49/100 x oil objective (Nikon Europe B.V., Amstelveen, Netherlands) together with Immersol™ 518F (Zeiss, Jena, Germany). Data was recorded by an Andor iXon 888 Ultra camera (Oxford Instruments, Tubney Woods, Abingdon, United Kingdom) equipped with an LS-OC optomask (Cairn Research Ltd, Faversham, United Kingdom) controlled with NIS-Elements AR 5.02.02. Single-molecule recordings were performed in a 256 × 256 pixel ROI in the camera's isolated cop mode. Movies of 10,000 frames (exposure time 10 ms, ≈88 fps, 115 s) of the cAMP signal were followed by movies of 3,000 frames (exposure time 40 ms; ≈25 fps; 75 s) of the eGFP s signal. Excitation and emission were filtered using appropriate filter sets (AHF Analysentechnik AG, Tübingen, Germany): eGFP-excitation: 472/30 Brighline HC, dichroic: zt 488 RDC, emission: 515/30 BrightLine HC (Semrock); f1cAMP: dichroic: ZT543rdc, emission: 596/83 BrightLine HC.

In widefield, eGFP was excited with a dimmed Nikon Intensilight lamp. For single molecule recording, eGFP was excited by a Stradus® 488-150 VORTRAN Laser (LASER

TECHNOLOGY, INC, City, USA/ Frankfurt Laser Company, Friedrichsdorf, Germany). Fluorescent f1cAMP derivatives were excited by a 100 mW MLL-III-543/1~100 mW laser (Changchun New Industries Optoelectronics Tech. Co., Changchun, China). Lasers were controlled by an AOTF included in the Nikon Laser Box LU4A.

**Single molecule analysis**

Single molecules were identified by iteratively selecting subsequent signal maxima in a maximal intensity projection of the background-subtracted movie and fitting a 2D Gaussian to 5 × 5 pixel around the maxima. Positions with fit results showing half axes wider than 2 pixels were rejected. The process was repeated at the same location for the eGFP-Signal. Due to the worse signal-to-noise ratio in the eGFP signal, leading to poorer fits, fits with half axes exceeding 3.3 pixels were rejected. If both the f1cAMP signal and the eGFP signal were present at the same position, the data trace was used for further analysis. The sum of a 3 × 3 area around the original location was used as signal for the traces. Traces were filtered by Chung-Kennedy filter with p, M and N being 40, 10 and [4 8 16 32][4]. Sparse and unusually bright one or two frame events were removed by a 5-point median filter.

From observed states the average probability of a ligand binding to a subunit, *P*(1), was calculated using

$$P(1) = \frac{N(1) + 2N(2) + 3N(3) + 4N(4)}{4(N(0) + N(1) + N(2) + N(3) + N(4))} \tag{1}$$

with N(x) being the number of frames spent in with x ligands bound. P(1) was used to calculate the expected binomial distribution according to:

$$F(x) = \binom{4}{x} P(1)^x \big(1 - P(1)\big)^{4-x} \tag{2}$$

with F(x) being the expected fraction of x Ligands bound.

Dwell times of single-liganded channels that were preceded and followed by states in which no ligand was bound were extracted and binned in a histogram. Mono- and biexponential distributions were fitted to the data. Functions follow the form of $Y = A_1 e^{-\frac{x}{\tau_1}}$ (monoexponential) and $Y = A_1 e^{-\frac{x}{\tau_1}} + A_2 e^{-\frac{x}{\tau_2}}$ (biexponential).

The quality of optimization was evaluated by ranking the maximum likelihood and relative errors of the parameters.

### Functional comparison of eGFPmHCN2 to mHCN2 channels

Wildtype mHCN2 and eGFPmHCN2 channels were expressed in *Xenopus laevis* oocytes. Patch-clamp recordings were performed in the inside-out configuration. Channel activation was studied by voltage jumps to -130 mV for both wildtype and eGFPmHCN2 channels (Extended Data Fig. 1 black curve). The cAMP-dependent modulation of channels was tested by application of a bath solution containing 10 µM cAMP. cAMP led to the expected increase in current in both wildtype and eGFPmHCN2 channels (Extended Data Fig. 1, red curve). For both channel types, activation, deactivation and amplitude increase were similar which confirms that the eGFPmHCN2 channels are suitable for the TIRF measurements.

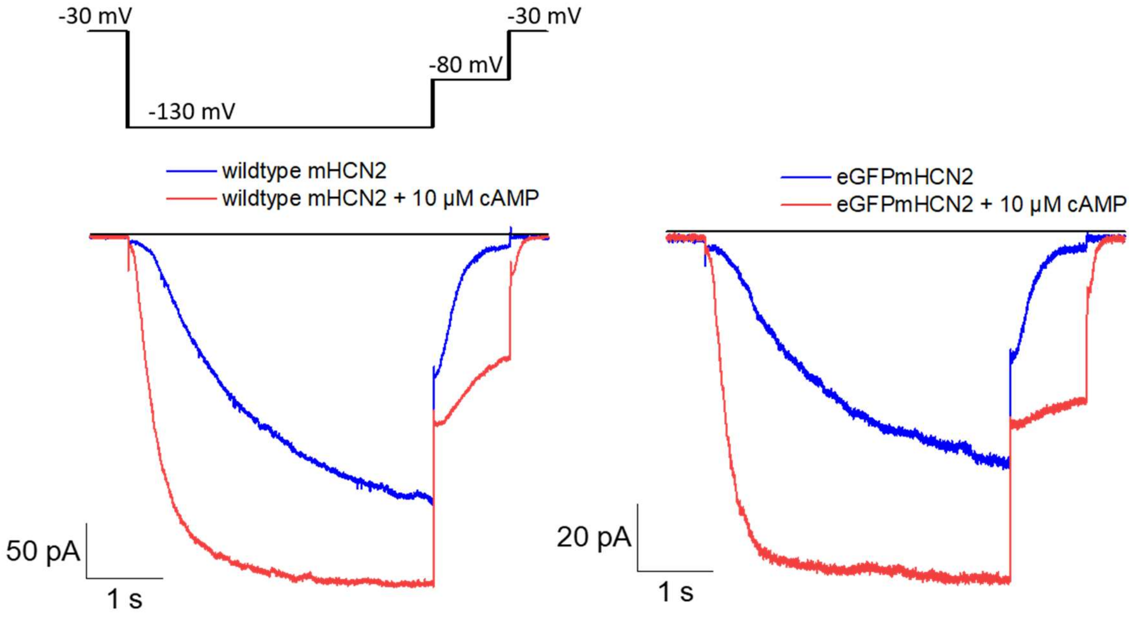


**Extended Data Fig.1. Comparison of the function of mHCN2 and eGFPmHCN2 channel.** The channels were expressed in *Xenopus* oocytes and the currents were measured in inside-out patches. In both cases 10 µM cAMP was applied.

### Channels are immobilized in native membranes

In the presented measurements, potential eGFP positions were pre-selected by calculating an image of maximal differences throughout the movie, filtering with PSF-sized Gaussian and subsequent selection of local maxima. These maxima were refined and filtered as channel positions by fitting the average of the first 100 images by the background-selected movie with a 2D-gaussian. Both photobleaching of individual GFP and ligand binding was evaluated by analyzing the sum of a 3 × 3 ROI centered at the fitted position.

We tested the mobility of eGFPmHCN2 channels by selecting a set of positions (N=9) with a point-spread function (PSF) like signals in both the GFP and ligand signal in the initial images and, subsequently, by tracking the eGFP signal. For this, a 2D Gaussian was fitted to a 7 × 7 pixel area centered with respect to the previous localization coordinates. From the fit coordinates, trajectories were plotted (Extended Data Fig. 2a) and a as function of time was calculated (Extended Data Fig. 2c). The latter converged rapidly to a maxima value, likely representing the special resolution at the given signal to noise. This interpretation was confirmed by a trajectory where the observed squared displacement increased with time (Extended Data Fig. 2c, black/grey curve: more detailed analysis showed that the increase coincided with GFP photobleaching (Extended Data Fig. 2b, blue curve), e.g. a decrease in the signal to noise ratio. Interestingly, the signal traces from the initial position showed systematically a more stable signal than traces constructed from the fitted position (Extended Data Fig. 2b, black/grey), confirming further that the fluctuation in the position is due to a deteriorating fit quality and not to an increased molecular mobility. In short, no indication of relevant eGFPmHCN2 mobility on the considered length scale was observed. We therefore did not see evidence arguing against the assumption of immobile receptors, which allowed us to use a small (3 × 3) ROI at a fixed position to follow ligand binding to eGFPmHCN2 channels. To rule out mechanical drifts, we tracked a very bright PSF-sized spot, likely an aggregate (Extended Data Fig. 2a, red). This showed almost no movement with the only deviations being single frame events that could arise from statistical artifacts. In summary, we saw no evidence arguing against the assumption of immobile receptors.

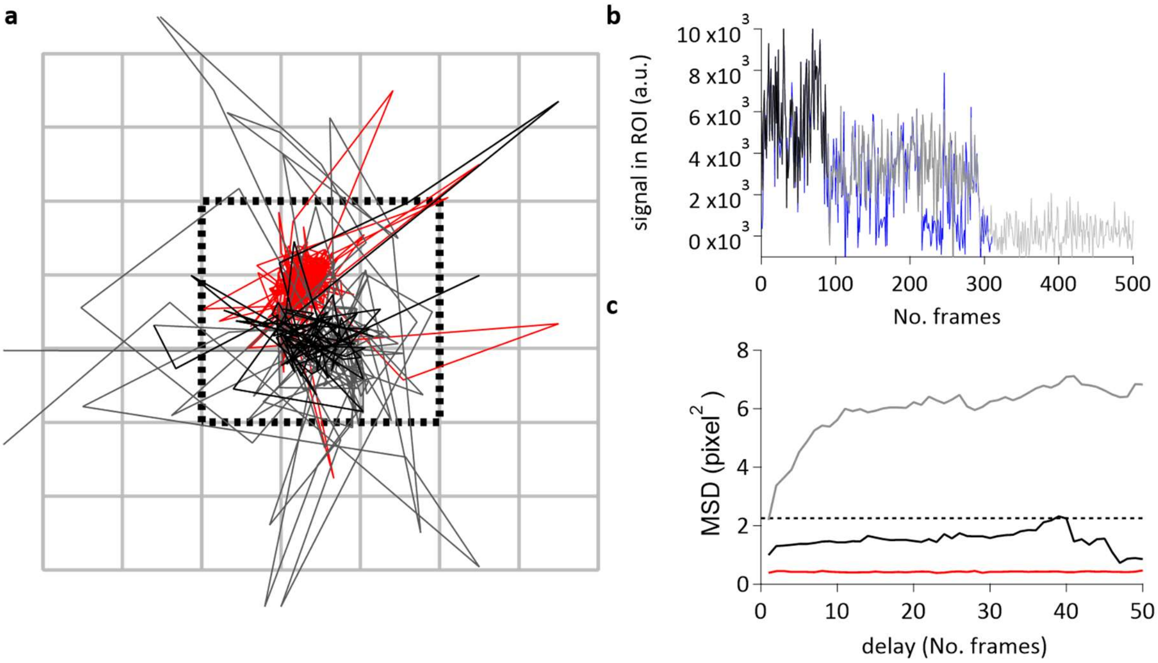


**Extended Data Fig. 2. eGFPmHCN2 channels are immobilized in native membranes. a,** Trajectory of an eGFPmHCN2 channel derived by tracking of eGFP signal with black to grey representing decreasing brightness. Red is a bright trajectory to test for mechanical stability. Grey squares show pixels (130 nm in sample space) and the dotted black line shows the ROI. **b,** eGFP-signal intensity of the tracked channel with black to grey corresponding to a. Intensities were calculated from the ROI at the initial position. In the blue curve the Intensity was calculated from a 3 × 3 ROI around the fit position for that frame. **c,** Mean square displacement of signal. Black dotted line indicates the displacement needed to leave the ROI. Black, grey, and red were calculated from the trajectories of the same color in a.

**Dwell time distribution of isolated single liganded states**

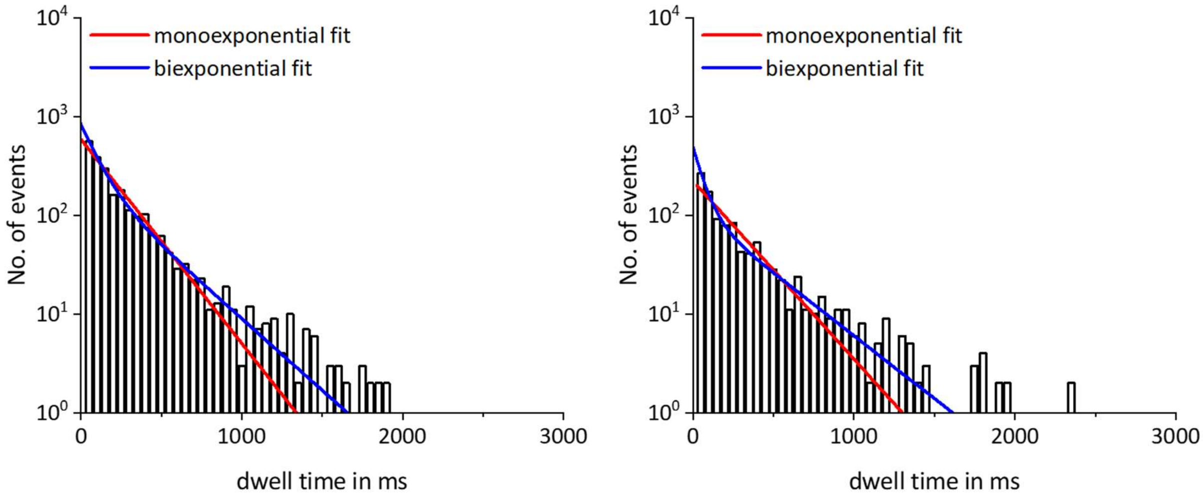


**Extended Data Fig. 3. Dwell time histograms of isolated single liganded states at different ligand concentrations.** Dwell times of single liganded states that were preceded and followed by empty states. Dwell times were recorded at 0.3 (left) and 0.5 µM f1cAMP (right). Either mono- or biexponential functions were fitted, suggesting a low and a high affinity state. In contrast, monoexponential fits fail to describe all data.

| Concentration of f1cAMP | Monoexponential fit | SEM | Biexponential fit | SEM |
|---|---|---|---|---|
| 0.1 | $A_1$: 510.7 | 10.4 | $A_1$: 476.5 | 18.9 |
| | $\tau_1$: 152.6 | 3.7 | $\tau_1$: 93.6 | 6.7 |
| | | | $A_2$: 121.2 | 24.0 |
| | | | $\tau_2$: 357.1 | 39.4 |
| 0.3 | $A_1$: 590.3 | 20.4 | $A_1$: 583.3 | 56.9 |
| | $\tau_1$: 209.9 | 5.1 | $\tau_1$: 97.0 | 11.5 |
| | | | $A_2$: 251.9 | 55.0 |
| | | | $\tau_2$: 299.8 | 23.1 |
| 0.5 | $A_1$: 222.6 | 11.9 | $A_1$: 383.7 | 67.5 |
| | $\tau_1$: 240.6 | 9.2 | $\tau_1$: 62.3 | 11.8 |
| | | | $A_2$: 113.1 | 16.0 |
| | | | $\tau_2$: 341.3 | 22.9 |

**Extended Data Table 1. Parameters describing mono and biexponential fit results from dwell time histograms.** The dimension of the time constants is ms.

### Model selection

A general assumption for our modeling was that ligand binding to one subunit has effects on the remaining empty subunits independent of their position (For models see Extended Data Table 1). To take into account both the tetrameric structure with four binding sites and the existence of a flip state, we first considered a model containing 10 states with up to four ligands bound in both the resting and the flip state (model 1). Out of the 26 rate constants, 4 are given due to microscopic reversibility, resulting in total in 22 free parameters for the rate constants. Models were analyzed using the QuB software. The log-likelihood (LL) value is provided by Extended Data Table 2.

Because in previous work models without binding in the flip states were reported[5], we also considered a respective model here (model 2). In our analysis, model 1 generated a higher LL than model 2 and it yielded binding and unbinding rate constants between the flip states that were in a similar range as in the resting states. Therefore, model 2 was discarded.

In the following, variations of model 1 are further considered. The main shortcoming for model 1 so far was that despite a reasonable convergence the overall parameter uncertainty was high. Therefore, the structure of model 1 was varied insofar that subsets of rate constants were equated to reduce the number of parameters. This is in contrast to model 1 where all rate constants were free parameters. In the resulting models 3 and 4, the binding rate constants in the resting, $k_x$, and the flip state, $k_x^*$, were equated, respectively. In model 5, the on-rate constants for flipping, $f_x$, were equated. For models 6,7 and 8, linear combinations of two of the restrictions used for model 3,4 and 5 were applied. For model 9, all three restrictions used for models 3, 4 and 5 were applied. The result was that that the LL became worse than that for model 1.

We next investigated models containing a flip state but without cooperativity, assuming either binding only in the resting state (model 10) or binding in both the resting and the flip state (model 11).

The models were ranked with the criterion of the higher LL, indicating a better fit. Generally, models with less parameters, e.g. by more equated rate constants, showed higher certainty of the estimated parameters though this led to a lower LL, i.e. a worse fit. Based on the LL, the non-cooperative model 10 would be best. However, this model contradicts the previously observed non-binomial distribution of observed states, since traces simulated from this model generate binomial distributions of observed states (Extended Data Fig. 3). The fit of model 11 did not converge. These two models were therefore not further considered.

The best model regarding the LL so far was model 1. However, the certainty of its parameters was too poor, to draw sufficiently substantiated conclusions on either cooperative or non-cooperative ligand binding. The second-best model with regard to the LL was model 7. Among all tested models with equated rate constants, the parameters had errors around 10%. However, this model shows extremely slow transition rates for ligand binding in the empty flipped channel and ligand unbinding in the single-liganded flipped channel. Therefore, we included model 13 which omits this transition. With respect to LL it was similar to model 7 while the certainty of the rate constants was further improved. Model 13 is therefore considered to be the best model for describing ligand binding in our experiments (Fig. 4a, b; Extended Data Table 2).

| Model-No. | Model structure | Rank | BIC/LL | No. of free Parameters |
|---|---|---|---|---|
| 1 | 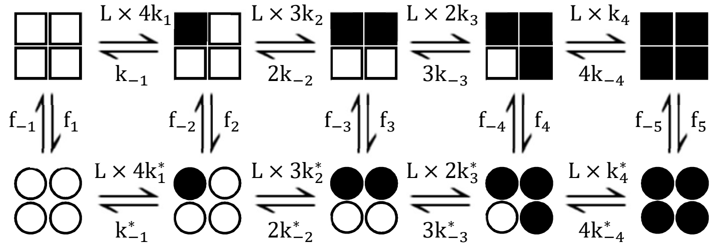  | 3 | -165723,67 | 22 |
| 2 | 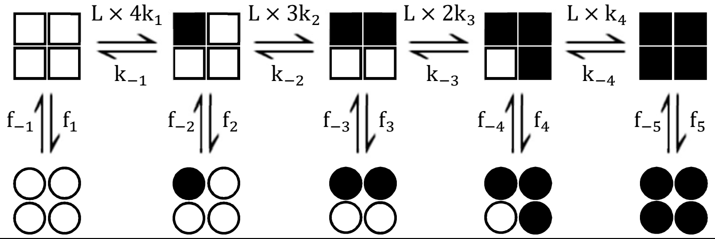  | 10 | -166190,32 | 18 |
| 3 | 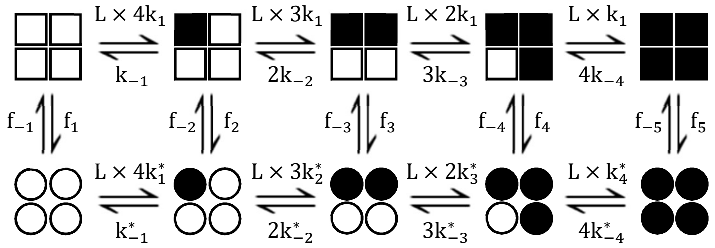  | 8 | -166597,98 | 19 |
| 4 | 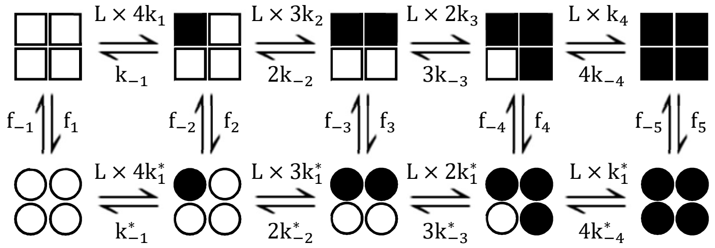  | 5 | -166190,32 | 19 |
| 5 | 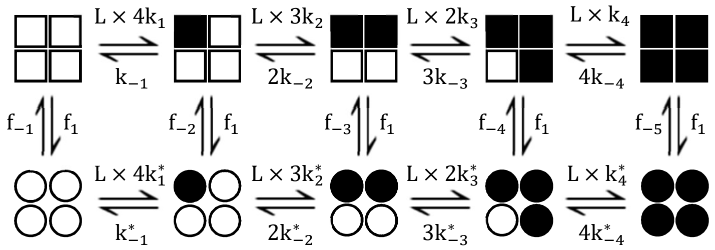  | 4 | -166126,77 | 18 |
| 6 | 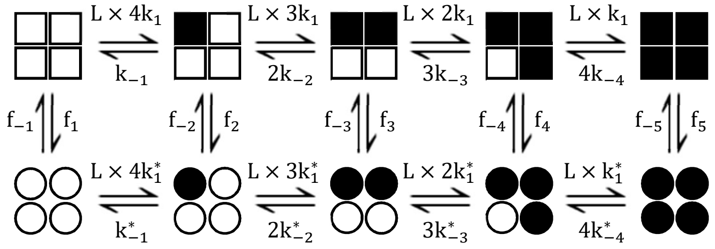  | 6 | -166399,38 | 19 |

| | | | | |
|---|---|---|---|---|
| 7 | 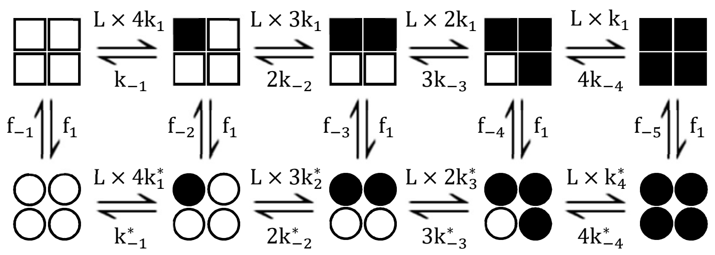 | 2 | -165961,90 | 15 |
| 8 | 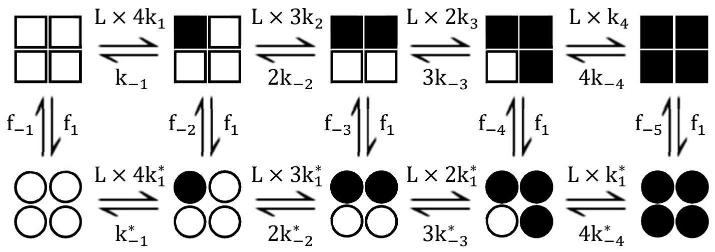 | 9 | -166751,94 | 15 |
| 9 | 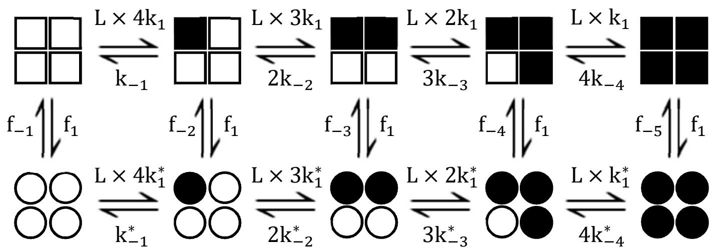 | 7 | -166421,05 | 12 |
| 10 | 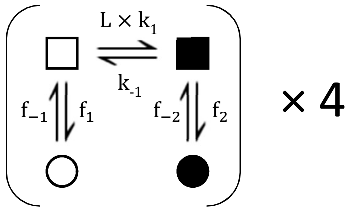 | 11 | 164318.55 | 6 |
| 11 | 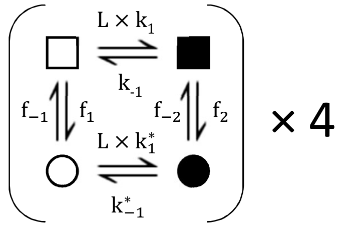 | 12 | No convergence | 7 |
| 12 | 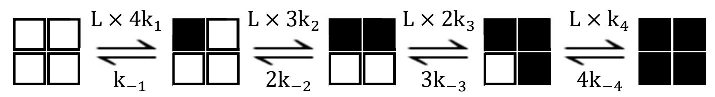 | 13 | Not valid see Fig. 3d | 8 |
| 13 | 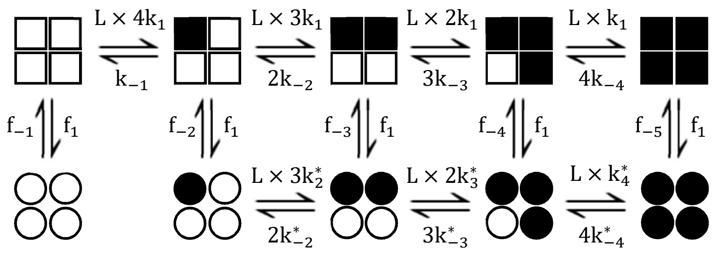 | 1 | -165961,90 | 14 |

**Extended Data Table 2. Models tested by maximum idealized point optimization.** Squares indicate that the subunit is in the resting state while circles show that the subunit is in a flip state. Open symbols indicate that a subunit is empty while filled symbols indicate an occupied subunit. L indicates the ligand concentration. Models were ranked based on the LL value with a higher number meaning a better fit and certainty of estimated parameters.

| Model No. | $k_{on}$ | value | %SEM | $k_{off}$ | value | %SEM | $K_X$ | value | %SEM |
|---|---|---|---|---|---|---|---|---|---|
| 1 | $k_1$ | $1.69\times10^{6}$ | 2.2% | $k_{-1}$ | $2.91\times10^{0}$ | 38.2% | $K_{A1}$ | $5.82\times10^{5}$ | 40.4% |
| | $k_2$ | $1.76\times10^{6}$ | 36.6% | $k_{-2}$ | $3.00\times10^{0}$ | 27.0% | $K_{A2}$ | $5.85\times10^{5}$ | 63.6% |
| | $k_3$ | $3.67\times10^{6}$ | 18.3% | $k_{-3}$ | $2.40\times10^{0}$ | 5.7% | $K_{A3}$ | $1.53\times10^{6}$ | 24.0% |
| | $k_4$ | $5.37\times10^{6}$ | 15.5% | $k_{-4}$ | $1.52\times10^{0}$ | 15.5% | $K_{A4}$ | $3.53\times10^{6}$ | 31.1% |
| | $k_1^*$ | $1.24\times10^{-1}$ | 68.8% | $k_{-1}^*$ | $1.80\times10^{-7}$ | 68.5% | $K_{A1}^*$ | $6.90\times10^{5}$ | 137.4% |
| | $k_2^*$ | $1.00\times10^{6}$ | 9.5% | $k_{-2}^*$ | $5.32\times10^{-1}$ | 9.5% | $K_{A2}^*$ | $1.88\times10^{6}$ | 18.9% |
| | $k_3^*$ | $1.00\times10^{6}$ | 5.5% | $k_{-3}^*$ | $6.19\times10^{-1}$ | 5.6% | $K_{A3}^*$ | $1.72\times10^{6}$ | 11.1% |
| | $k_4^*$ | $1.71\times10^{6}$ | 15.2% | $k_{-4}^*$ | $5.23\times10^{-1}$ | 15.2% | $K_{A4}^*$ | $3.28\times10^{6}$ | 30.4% |
| | $f_1$ | 0.489 | 16.7% | $f_{-1}$ | 0.635 | 15.9% | $K_{F1}$ | 0.770 | 32.6% |
| | $f_2$ | 0.530 | 10.9% | $f_{-2}$ | 0.581 | 10.1% | $K_{F2}$ | 0.912 | 21.0% |
| | $f_3$ | 1.299 | 13.4% | $f_{-3}$ | 0.443 | 12.3% | $K_{F3}$ | 2.933 | 25.6% |
| | $f_4$ | 0.465 | 26.5% | $f_{-4}$ | 0.141 | 25.7% | $K_{F4}$ | 3.285 | 52.1% |
| | $f_5$ | 0.137 | 65.1% | $f_{-5}$ | 0.045 | 63.4% | $K_{F5}$ | 3.053 | 128.5% |
| 2 | $k_1$ | $6.76\times10^{6}$ | 2.2% | $k_{-1}$ | $2.53\times10^{0}$ | 2.1% | $K_{A1}$ | $2.67\times10^{6}$ | 4.4% |
| | $k_2$ | $7.07\times10^{6}$ | 2.3% | $k_{-2}$ | $4.12\times10^{0}$ | 2.6% | $K_{A2}$ | $1.71\times10^{6}$ | 4.9% |
| | $k_3$ | $6.03\times10^{6}$ | 3.0% | $k_{-3}$ | $4.88\times10^{0}$ | 3.2% | $K_{A3}$ | $1.24\times10^{6}$ | 6.2% |
| | $k_4$ | $3.98\times10^{6}$ | 4.5% | $k_{-4}$ | $4.75\times10^{0}$ | 5.1% | $K_{A4}$ | $8.38\times10^{6}$ | 9.6% |
| | $f_1$ | 0.488 | 7.9% | $f_{-1}$ | 0.635 | 5.0% | $K_{F1}$ | 0.769 | 13.0% |
| | $f_2$ | 1.293 | 9.6% | $f_{-2}$ | 1.923 | 5.8% | $K_{F2}$ | 0.672 | 15.4% |
| | $f_3$ | 1.613 | 10.6% | $f_{-3}$ | 2.102 | 6.5% | $K_{F3}$ | 0.768 | 17.1% |
| | $f_4$ | 1.081 | 15.4% | $f_{-4}$ | 1.901 | 8.8% | $K_{F4}$ | 0.569 | 24.1% |
| | $f_5$ | 0.624 | 23.9% | $f_{-5}$ | 1.221 | 16.1% | $K_{F5}$ | 0.512 | 40.0% |
| 3 | $k_1$ | $1.32\times10^{6}$ | 2.0% | $k_{-1}$ | $3.50\times10^{-2}$ | 22.6% | $K_{A1}$ | $3.78\times10^{6}$ | 24.6% |
| | | | | $k_{-2}$ | $7.43\times10^{-1}$ | 38.1% | $K_{A2}$ | $1.78\times10^{6}$ | 40.1% |
| | | | | $k_{-3}$ | $7.40\times10^{-1}$ | 28.2% | $K_{A3}$ | $1.78\times10^{6}$ | 30.2% |
| | | | | $k_{-4}$ | $5.38\times10^{-1}$ | 35.9% | $K_{A4}$ | $2.46\times10^{6}$ | 38.0% |
| | $k_1^*$ | $8.17\times10^{5}$ | 16.0% | $k_{-1}^*$ | $3.59\times10^{0}$ | 16.1% | $K_{A1}^*$ | $2.28\times10^{5}$ | 32.1% |
| | $k_2^*$ | $1.79\times10^{6}$ | 6.7% | $k_{-2}^*$ | $3.52\times10^{0}$ | 7.3% | $K_{A2}^*$ | $5.10\times10^{5}$ | 14.0% |
| | $k_3^*$ | $4.66\times10^{6}$ | 11.0% | $k_{-3}^*$ | $2.62\times10^{0}$ | 10.9% | $K_{A3}^*$ | $1.78\times10^{6}$ | 21.9% |
| | $k_4^*$ | $7.75\times10^{6}$ | 18.0% | $k_{-4}^*$ | $1.54\times10^{0}$ | 18.2% | $K_{A4}^*$ | $5.02\times10^{6}$ | 36.1% |
| | $f_1$ | $4.35\times10^{-8}$ | 61.0% | $f_{-1}$ | $4.47\times10^{-9}$ | 61.0% | $K_{F1}$ | 9.727 | 122.0% |
| | $f_2$ | 0.234 | 15.7% | $f_{-2}$ | 0.398 | 15.3% | $K_{F2}$ | 0.587 | 31.0% |
| | $f_3$ | 0.251 | 13.5% | $f_{-3}$ | 1.490 | 13.7% | $K_{F3}$ | 0.168 | 27.2% |
| | $f_4$ | 0.151 | 32.5% | $f_{-4}$ | 0.899 | 32.1% | $K_{F4}$ | 0.168 | 64.6% |
| | $f_5$ | 0.057 | 60.7% | $f_{-5}$ | 0.165 | 63.1% | $K_{F5}$ | 0.344 | 123.8% |
| 4 | $k_1$ | $1.69\times10^{6}$ | 2.1% | $k_{-1}$ | $2.53\times10^{0}$ | 25.5% | $K_{A1}$ | $6.69\times10^{5}$ | 27.6% |
| | $k_2$ | $2.36\times10^{6}$ | 26.4% | $k_{-2}$ | $2.06\times10^{0}$ | 29.8% | $K_{A2}$ | $1.14\times10^{6}$ | 56.2% |
| | $k_3$ | $3.01\times10^{6}$ | 19.2% | $k_{-3}$ | $1.63\times10^{0}$ | 27.7% | $K_{A3}$ | $1.85\times10^{6}$ | 46.8% |
| | $k_4$ | $3.98\times10^{6}$ | 28.3% | $k_{-4}$ | $1.19\times10^{0}$ | 27.5% | $K_{A4}$ | $3.35\times10^{6}$ | 55.9% |
| | $k_1^*$ | $5.22\times10^{-3}$ | 15.6% | $k_{-1}^*$ | $8.92\times10^{-9}$ | 30.6% | $K_{A1}^*$ | $5.85\times10^{5}$ | 46.2% |
| | | | | $k_{-2}^*$ | $4.00\times10^{-9}$ | 32.2% | $K_{A2}^*$ | $1.30\times10^{6}$ | 47.8% |

| | | | | | | | | | |
|---|---|---|---|---|---|---|---|---|---|
| | | | | $k^*_{-3}$ | $3.80\times10^{-9}$ | 32.4% | $K^*_{A3}$ | $1.37\times10^{6}$ | 48.0% |
| | | | | $k^*_{-4}$ | $1.73\times10^{-9}$ | 34.9% | $K^*_{A4}$ | $3.01\times10^{6}$ | 50.5% |
| | $f_1$ | 0.488 | 9.4% | $f_{-1}$ | 0.635 | 8.3% | $K_{F1}$ | 0.769 | 17.7% |
| | $f_2$ | 1.293 | 11.7% | $f_{-2}$ | 1.923 | 10.3% | $K_{F2}$ | 0.672 | 22.1% |
| | $f_3$ | 1.613 | 11.6% | $f_{-3}$ | 2.102 | 9.7% | $K_{F3}$ | 0.768 | 21.3% |
| | $f_4$ | 1.081 | 16.1% | $f_{-4}$ | 1.901 | 14.1% | $K_{F4}$ | 0.569 | 30.3% |
| | $f_5$ | 0.624 | 22.4% | $f_{-5}$ | 1.221 | 15.7% | $K_{F5}$ | 0.512 | 38.1% |
| 5 | $k_1$ | $4.66\times10^{6}$ | 4.2% | $k_{-1}$ | $5.02\times10^{-1}$ | 5.9% | $K_{A1}$ | $9.28\times10^{6}$ | 10.2% |
| | $k_2$ | $1.18\times10^{6}$ | 6.3% | $k_{-2}$ | $7.89\times10^{-1}$ | 6.2% | $K_{A2}$ | $1.49\times10^{6}$ | 12.4% |
| | $k_3$ | $1.20\times10^{6}$ | 6.9% | $k_{-3}$ | $7.41\times10^{-1}$ | 6.4% | $K_{A3}$ | $1.62\times10^{6}$ | 13.3% |
| | $k_4$ | $1.65\times10^{6}$ | 7.9% | $k_{-4}$ | $5.42\times10^{-1}$ | 7.4% | $K_{A4}$ | $3.05\times10^{6}$ | 15.3% |
| | $k^*_1$ | $7.28\times10^{5}$ | 3.2% | $k^*_{-1}$ | $3.99\times10^{0}$ | 3.4% | $K^*_{A1}$ | $1.83\times10^{5}$ | 6.6% |
| | $k^*_2$ | $1.89\times10^{6}$ | 3.0% | $k^*_{-2}$ | $3.71\times10^{0}$ | 3.6% | $K^*_{A2}$ | $5.08\times10^{5}$ | 6.6% |
| | $k^*_3$ | $5.41\times10^{6}$ | 4.8% | $k^*_{-3}$ | $2.76\times10^{0}$ | 4.8% | $K^*_{A3}$ | $1.96\times10^{6}$ | 9.6% |
| | $k^*_4$ | $8.16\times10^{6}$ | 7.0% | $k^*_{-4}$ | $1.66\times10^{0}$ | 7.8% | $K^*_{A4}$ | $4.93\times10^{6}$ | 14.9% |
| | $f_1$ | 0.238 | 3.3% | $f_{-1}$ | 0.011 | 7.8% | $K_{F1}$ | 20.836 | 11.1% |
| | | | | $f_{-2}$ | 0.582 | 5.9% | $K_{F2}$ | 0.410 | 9.2% |
| | | | | $f_{-3}$ | 1.711 | 6.1% | $K_{F3}$ | 0.139 | 9.3% |
| | | | | $f_{-4}$ | 1.413 | 8.1% | $K_{F4}$ | 0.169 | 11.3% |
| | | | | $f_{-5}$ | 0.874 | 13.8% | $K_{F5}$ | 0.273 | 17.0% |
| 6 | $k_1$ | $2.33\times10^{6}$ | 1.3% | $k_{-1}$ | $2.53\times10^{0}$ | 28.3% | $K_{A1}$ | $9.22\times10^{5}$ | 29.6% |
| | | | | $k_{-2}$ | $1.83\times10^{0}$ | 28.6% | $K_{A2}$ | $1.28\times10^{6}$ | 29.9% |
| | | | | $k_{-3}$ | $1.42\times10^{0}$ | 29.7% | $K_{A3}$ | $1.64\times10^{6}$ | 31.1% |
| | | | | $k_{-4}$ | $1.16\times10^{0}$ | 28.9% | $K_{A4}$ | $2.01\times10^{6}$ | 30.2% |
| | $k^*_1$ | $3.98\times10^{-3}$ | 15.2% | $k^*_{-1}$ | $8.05\times10^{-9}$ | 32.0% | $K^*_{A1}$ | $4.94\times10^{5}$ | 47.2% |
| | | | | $k^*_{-2}$ | $3.65\times10^{-9}$ | 31.5% | $K^*_{A2}$ | $1.09\times10^{6}$ | 46.7% |
| | | | | $k^*_{-3}$ | $3.62\times10^{-9}$ | 31.6% | $K^*_{A3}$ | $1.10\times10^{6}$ | 46.8% |
| | | | | $k^*_{-4}$ | $1.52\times10^{-9}$ | 34.7% | $K^*_{A4}$ | $2.62\times10^{6}$ | 49.9% |
| | $f_1$ | 1.090 | 25.1% | $f_{-1}$ | 0.877 | 24.8% | $K_{F1}$ | | |
| | $f_2$ | 1.266 | 24.3% | $f_{-2}$ | 1.901 | 13.4% | $K_{F2}$ | 1.242 | 49.9% |
| | $f_3$ | 0.898 | 12.5% | $f_{-3}$ | 1.581 | 11.4% | $K_{F3}$ | 0.666 | 37.7% |
| | $f_4$ | 0.514 | 15.6% | $f_{-4}$ | 1.349 | 13.6% | $K_{F4}$ | 0.568 | 23.9% |
| | $f_5$ | 0.598 | 22.2% | $f_{-5}$ | 1.201 | 16.8% | $K_{F5}$ | 0.381 | 29.2% |
| | | | | | | | | 0.498 | 39.0% |
| 7 | $k_1$ | $1.92\times10^{6}$ | 1.5% | $k_{-1}$ | $2.91\times10^{0}$ | 37.7% | $K_{A1}$ | $6.59\times10^{5}$ | 39.3% |
| | | | | $k_{-2}$ | $2.75\times10^{0}$ | 14.9% | $K_{A2}$ | $6.98\times10^{5}$ | 16.4% |
| | | | | $k_{-3}$ | $2.42\times10^{0}$ | 14.7% | $K_{A3}$ | $7.92\times10^{5}$ | 16.3% |
| | | | | $k_{-4}$ | $1.70\times10^{0}$ | 22.6% | $K_{A4}$ | $1.13\times10^{6}$ | 24.1% |
| | $k^*_1$ | $2.58\times10^{-4}$ | 59.4% | $k^*_{-1}$ | $3.88\times10^{-10}$ | 59.4% | $K^*_{A1}$ | $6.65\times10^{5}$ | 118.8% |
| | $k^*_2$ | $8.93\times10^{5}$ | 6.6% | $k^*_{-2}$ | $5.02\times10^{-1}$ | 6.6% | $K^*_{A2}$ | $1.78\times10^{6}$ | 13.2% |
| | $k^*_3$ | $1.30\times10^{6}$ | 5.4% | $k^*_{-3}$ | $6.88\times10^{-1}$ | 5.3% | $K^*_{A3}$ | $1.89\times10^{6}$ | 10.8% |
| | $k^*_4$ | $2.20\times10^{6}$ | 8.7% | $k^*_{-4}$ | $6.17\times10^{-1}$ | 8.6% | $K^*_{A4}$ | $3.57\times10^{6}$ | 17.3% |
| | $f_1$ | 0.621 | 4.1% | $f_{-1}$ | 0.687 | 21.9% | $K_{F1}$ | 0.904 | 25.9% |
| | | | | $f_{-2}$ | 0.681 | 7.8% | $K_{F2}$ | 0.912 | 11.8% |
| | | | | $f_{-3}$ | 0.267 | 10.5% | $K_{F3}$ | 2.324 | 14.6% |

| | | | | | | | | | |
|---|---|---|---|---|---|---|---|---|---|
| | | | | $f_{-4}$ | 0.112 | 7.4% | $K_{F4}$ | 5.549 | 11.5% |
| | | | | $f_{-5}$ | 0.035 | 13.9% | $K_{F5}$ | 17.538 | 17.9% |
| 8 | $k_1$ | $7.93\times10^{5}$ | 1.9% | $k_{-1}$ | $3.78\times10^{0}$ | 7.7% | $K_{A1}$ | $2.13\times10^{5}$ | 9.6% |
| | $k_2$ | $1.68\times10^{6}$ | 10.0% | $k_{-2}$ | $3.28\times10^{0}$ | 8.3% | $K_{A2}$ | $5.13\times10^{5}$ | 18.2% |
| | $k_3$ | $4.00\times10^{6}$ | 9.1% | $k_{-3}$ | $2.29\times10^{0}$ | 12.0% | $K_{A3}$ | $1.75\times10^{6}$ | 21.2% |
| | $k_4$ | $6.58\times10^{6}$ | 10.3% | $k_{-4}$ | $1.48\times10^{0}$ | 9.8% | $K_{A4}$ | $4.46\times10^{6}$ | 20.0% |
| | $k_1^*$ | $1.24\times10^{-1}$ | 3.5% | $k_{-1}^*$ | $4.61\times10^{-1}$ | 7.3% | $K_{A1}^*$ | $3.05\times10^{6}$ | 10.8% |
| | | | | $k_{-2}^*$ | $8.20\times10^{-1}$ | 3.5% | $K_{A2}^*$ | $1.72\times10^{6}$ | 7.0% |
| | | | | $k_{-3}^*$ | $8.19\times10^{-1}$ | 4.9% | $K_{A3}^*$ | $1.72\times10^{6}$ | 8.4% |
| | | | | $k_{-4}^*$ | $6.03\times10^{-1}$ | 7.0% | $K_{A4}^*$ | $2.33\times10^{6}$ | 10.5% |
| | $f_1$ | 0.188 | 6.2% | $f_{-1}$ | 1.577 | 7.4% | $K_{F1}$ | 0.119 | 13.6% |
| | | | | $f_{-2}$ | 0.110 | 7.1% | $K_{F2}$ | 1.709 | 13.3% |
| | | | | $f_{-3}$ | 0.033 | 8.9% | $K_{F3}$ | 5.715 | 15.0% |
| | | | | $f_{-4}$ | 0.033 | 13.1% | $K_{F4}$ | 5.628 | 19.2% |
| | | | | $f_{-5}$ | 0.064 | 15.8% | $K_{F5}$ | 2.948 | 22.0% |
| 9 | $k_1$ | $2.28\times10^{6}$ | 1.4% | $k_{-1}$ | $2.43\times10^{0}$ | 25.1% | $K_{A1}$ | $9.36\times10^{5}$ | 26.4% |
| | | | | $k_{-2}$ | $1.84\times10^{0}$ | 27.8% | $K_{A2}$ | $1.24\times10^{6}$ | 29.1% |
| | | | | $k_{-3}$ | $1.53\times10^{0}$ | 30.3% | $K_{A3}$ | $1.49\times10^{6}$ | 31.7% |
| | | | | $k_{-4}$ | $1.23\times10^{0}$ | 19.6% | $K_{A4}$ | $1.85\times10^{6}$ | 20.9% |
| | $k_1^*$ | $1.07\times10^{-3}$ | 15.1% | $k_{-1}^*$ | $2.27\times10^{-9}$ | 30.9% | $K_{A1}^*$ | $4.72\times10^{5}$ | 45.9% |
| | | | | $k_{-2}^*$ | $8.72\times10^{-10}$ | 30.9% | $K_{A2}^*$ | $1.23\times10^{6}$ | 46.0% |
| | | | | $k_{-3}^*$ | $8.16\times10^{-10}$ | 29.8% | $K_{A3}^*$ | $1.31\times10^{6}$ | 44.9% |
| | | | | $k_{-4}^*$ | $4.67\times10^{-10}$ | 30.6% | $K_{A4}^*$ | $2.29\times10^{6}$ | 45.7% |
| | $f_1$ | 0.977 | 14.4% | $f_{-1}$ | 0.828 | 21.7% | $K_{F1}$ | 1.180 | 36.1% |
| | | | | $f_{-2}$ | 1.645 | 26.8% | $K_{F2}$ | 0.594 | 41.1% |
| | | | | $f_{-3}$ | 1.661 | 25.6% | $K_{F3}$ | 0.588 | 40.0% |
| | | | | $f_{-4}$ | 1.892 | 14.6% | $K_{F4}$ | 0.517 | 28.9% |
| | | | | $f_{-5}$ | 1.527 | 12.5% | $K_{F5}$ | 0.640 | 26.8% |
| 10 | $k_1$ | $2.60\times10^{6}$ | 2.2% | $k_{-1}$ | $2.76\times10^{0}$ | 4.8% | $K_{A1}$ | $9.43\times10^{5}$ | 7.0% |
| | $f_1$ | 0.099 | 5.8% | $f_{-1}$ | 0.092 | 5.7% | $K_{F1}$ | 1.073 | 11.4% |
| | $f_2$ | 0.473 | 6.7% | $f_{-2}$ | 0.357 | 4.9% | $K_{F2}$ | 1.324 | 11.7% |
| 11 | | Did not converge | | | | | | | |
| 12 | $k_1$ | $3.82\times10^{6}$ | 1.4% | $k_{-1}$ | $1.37\times10^{0}$ | 1.5% | $K_{A1}$ | $2.79\times10^{6}$ | 2.9% |
| | $k_2$ | $3.88\times10^{6}$ | 1.6% | $k_{-2}$ | $2.23\times10^{0}$ | 1.6% | $K_{A2}$ | $1.74\times10^{6}$ | 3.2% |
| | $k_3$ | $3.29\times10^{6}$ | 2.2% | $k_{-3}$ | $3.04\times10^{0}$ | 2.1% | $K_{A3}$ | $1.08\times10^{6}$ | 4.3% |
| | $k_4$ | $2.61\times10^{6}$ | 4.1% | $k_{-4}$ | $3.11\times10^{0}$ | 4.6% | $K_{A4}$ | $8.39\times10^{5}$ | 8.7% |

| | | | | | | | | | |
|---|---|---|---|---|---|---|---|---|---|
| 13 | $k_1$ | $1.92\times10^6$ | 1.5% | $k_{-1}$ | $2.91\times10^0$ | 5.3% | $K_{A1}$ | $6.59\times10^5$ | 6.8% |
| | | | | $k_{-2}$ | $2.75\times10^0$ | 6.0% | $K_{A2}$ | $6.98\times10^5$ | 7.5% |
| | | | | $k_{-3}$ | $2.42\times10^0$ | 5.6% | $K_{A3}$ | $7.92\times10^5$ | 7.1% |
| | | | | $k_{-4}$ | $1.70\times10^0$ | 8.9% | $K_{A4}$ | $1.13\times10^6$ | 10.5% |
| | $k_2^*$ | $8.93\times10^5$ | 4.2% | $k_{-2}^*$ | $5.02\times10^{-1}$ | 3.7% | $K_{A2}^*$ | $1.78\times10^6$ | 7.9% |
| | $k_3^*$ | $1.30\times10^6$ | 3.4% | $k_{-3}^*$ | $6.88\times10^{-1}$ | 3.7% | $K_{A3}^*$ | $1.89\times10^6$ | 7.0% |
| | $k_4^*$ | $2.20\times10^6$ | 5.6% | $k_{-4}^*$ | $6.17\times10^{-1}$ | 5.1% | $K_{A4}^*$ | $3.57\times10^6$ | 10.7% |
| | $f_1$ | 0.621 | 3.0% | $f_{-1}$ | 0.687 | 6.9% | $K_{F1}$ | 0.621 | 9.9% |
| | | | | $f_{-2}$ | 0.681 | 4.4% | $K_{F2}$ | 0.912 | 7.4% |
| | | | | $f_{-3}$ | 0.267 | 5.0% | $K_{F3}$ | 2.324 | 8.0% |
| | | | | $f_{-4}$ | 0.112 | 8.3% | $K_{F4}$ | 5.549 | 11.4% |
| | | | | $f_{-5}$ | 0.035 | 13.9% | $K_{F5}$ | 17.538 | 17.0% |

**Extended data table 3. Constants for the models.** Association rate constants ($k_{on}$, in $M^{-1}s^{-1}$), dissociation rate constants ($k_{off}$, in $s^{-1}$), equilibrium association constants ($K_A$, in $M^{-1}$), flip rate constants ($k_{flip}$, in $s^{-1}$; $k_{-flip}$, in $s^{-1}$), and flip equilibrium constants ($K_{flip}$, dimensionless) of tested models.

### Supplementary Discussion

While model 10 has a lower LL value than all other models tested and therefore seems to provide a better fit, it contradicts earlier observations. The distribution of states observed differed significantly from a binomial model. The transition rates from model 10 were used to simulate as many data points and traces over the same concentration range as was used for estimation of the parameters. These simulated data traces were analyzed for distribution of the observed states. The distribution was compared to a binomial distribution and matched the expected ratios. No significant difference was found. From this, we conclude that the lower LL does not come from a better fit of the data, but the lower number of parameters.

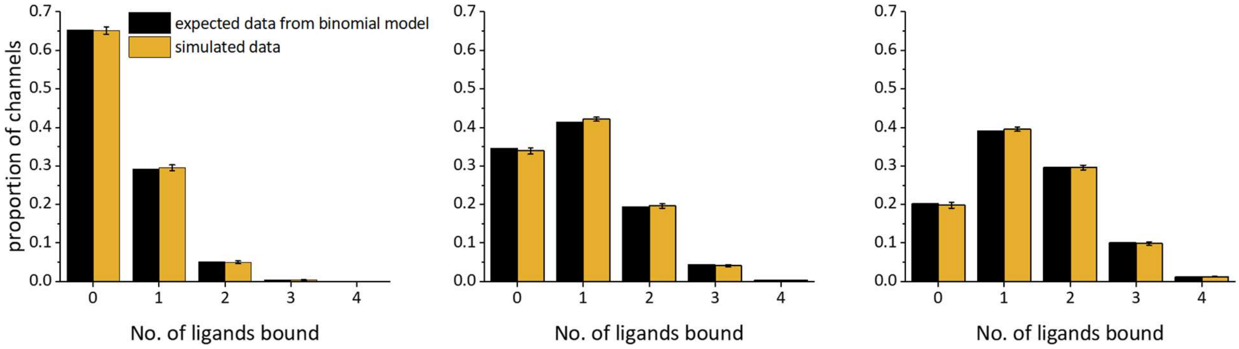


**Extended Data Fig. 4. Comparison of observed and expected distribution of simulated traces from a non-cooperative model.** Comparison of state distribution of data simulated with model 10 to the expected distribution for a binomial model for the f1cAMP concentrations of 0.1 – 0.5 µM. There were no significant differences (one sample *t*-test, $P<0.05$).

| No. of ligands bound | 0.1 µM fcAMP | | 0.3 µM fcAMP | | 0.5 µM fcAMP | |
|---|---|---|---|---|---|---|
| | expected portion | observed portion | expected portion | observed portion | expected portion | observed portion |
| 0 | 65.12 % | 65.28 ± 0.96 % | 33.88 % | 34.49 ± 0.81 % | 19.79 % | 20.11 ± 0.74 % |
| 1 | 29.48 % | 29.19 ± 0.76 % | 42.11 % | 41.38 ± 0.52 % | 39.52 % | 39.05 ± 0.58 % |
| 2 | 5.01 % | 5.14 ± 0.35 % | 19.63 % | 19.36 ± 0.57 % | 29.6 % | 29.59 ± 0.62 % |
| 3 | 0.38 % | 0.37 ± 0.05 % | 4.07 % | 4.36 ± 0.29 % | 9.85 % | 10.03 ± 0.50 % |
| 4 | 0.01 % | 0.02 ± 0.01 % | 0.32 % | 0.41 ± 0.06 % | 1.23 % | 1.22 ± 0.13 % |

**Extended Data Table 4. Observed and expected portions of liganded states from simulated data.**

### Suppression of f1cAMP binding to endogenous membrane proteins

Since we worked with membranes originating from functional cells, there are also endogenous proteins in membrane. Using the harmonizome database, we took into account cAMP binding membrane proteins that are potentially expressed endogenously by HEK293 cells[6]. Apart from mHCN2, we identified as potential cAMP, and thus f1cAMP, binding proteins in HEK293 cells protein kinase A (PKA), adenylate cyclase (AC), phosphodiesterase (PDE) or exchange Protein activated by cAMP (EPAC). Binding of f1cAMP to one of those endogenous proteins would result in false positive binding signals if they were in close proximity of a fluorescently labeled mHCN2 channel. To reduce this potential background binding, we searched for tested selective competitive inhibitors that do not bind to and affect mHCN2 channels (Extended Data Fig 5b). The following substances were identified from literature: Binding to AC can be neglected since this would require initial binding of pyrophosphate which is not present in the applied solutions[7]. Binding to PDE can be prevented by applying saturating concentrations of Piclamilast[8]. Binding to EPAC can be blocked by the modified cAMP analogue 8-pCPT-2-OMe-cAMP[9]. For the PKA we did not find any selective competitive inhibitor that would also not bind to mHCN2. Because of this we used ht31, an A – Kinase anchor protein analogue[10]. This peptide competes for the anchor that attaches the regulatory subunit to the membrane. By adding sufficient ht31, we assumed to replace the bound PKA with the anchor protein analogue, thus detaching the regulatory subunit from the membrane and removing it by solution exchange.

To evaluate the potency of these competitive inhibitors, we evaluated the binding of f1cAMP to the native, not HCN2-expressing, membrane. Membranes were identified by staining them with the membrane dye DiO (Extended Data Fig. 4a (I)). We then identified binding events inside the membrane area with the algorithm described in Methods. The binding event density included f1cAMP binding to endogenous cAMP binding membrane proteins, but also f1cAMP interacting non-specifically with the membrane surface. These random movements would also be counted as binding to the membrane. To get an estimate of these random binding events, we measured the density of binding events when 0.1 µM f1cAMP was added to the bath solution and compared this density to another measurement where 1 mM cAMP was added to compete with a specific binding (Extended Data Fig. 5c). When no cAMP was present, we observed an average density of binding events of 1.33 ± 0.01 $\mu m^{-2}$. When 1 mM of cAMP was added the average density of binding events decreased to 0.98 ± 0.01 $\mu m^{-2}$. This means that on average there are 0.35 ± 0.02 f1cAMP binding molecules per $\mu m^2$ of membrane, while 0.98 ± 0.01 binding events per $\mu m^2$ appeared most likely due to random interactions or pure diffusion on and off the surface.

In the next step, we investigated the potency of our tested inhibitor mixture. We repeated the experiment with the selective competitive inhibitors instead of 1 mM cAMP. While the competitive inhibitors were present, the average density of binding events decreased from 1.33 ± 0.01 $\mu m^{-2}$ to 1.01 ± 0.01 $\mu m^{-2}$. This means that the mixture of inhibitors can suppress binding to about 0.32 ± 0.01 f1cAMP binding proteins per $\mu m^2$. Thus, the inhibitor mixture suppresses up to 92 % of specific endogenous binding sites.

This percentage was the maximum of inhibition that we could achieve with the already mentioned components. Therefore, it is likely that there are other membrane proteins that can also bind f1cAMP and cAMP that were not targeted by the inhibitors mentioned here. Examples could be cGMP dependent proteins since cGMP and cAMP have lots of structural similarities and there are cGMP binders known to also bind cAMP[11]. Another possible explanation could be that there are cAMP binding molecules that we do not know yet or are not well enough studied so far to identify competitive inhibitors, such as the popeye domain[12]. However, since we could already block more than 90 % of the endogenous signal, these other binding proteins were neglected in the analysis.

We still needed to ensure that the inhibitors did not interfere with the binding of cAMP to mHCN2 channels. We therefore performed patch-clamp measurements. mHCN2 channels were expressed in *Xenopus laevis* oocytes and the channels were investigated in patch-clamped macropatches in the inside-out configuration. The patches were kept at -30 mV and the HCN-channel current was evoked by stepping to -130 mV. After 5 s the voltage was again stepped to -30 mV. The experiment was then repeated in the presence of 0.1 µM cAMP, a concentration near the EC50 value for mHCN2 channels[13]. To test the inhibitors, the patches were then exposed to 0.1 µM cAMP plus the inhibitor concentrations used in the TIRF experiments. The system was equilibrated for 5 minutes. Since we applied cAMP at the EC50 concentration there should be an increase in current if any of the inhibitors would bind and activate the channel. If any of the inhibitors would bind but not activate the channel, we would see a decrease in current because the binding sites would get blocked for cAMP and therefore the channels would be less activated. We did neither observe a change in activation nor deactivation kinetics nor current amplitude when the inhibitors were applied (Extended Data Fig. 5d). To verify that the channel functionality is also kept after the application, we washed the inhibitors off and measured channel properties again with only cAMP added, as well as only bath medium. The activation curves match each other almost perfectly. Overall, we conclude that the mixture of inhibitors does not relevantly interact with the mHCN2 channel binding site and is therefore suited for the TIRF binding experiments.

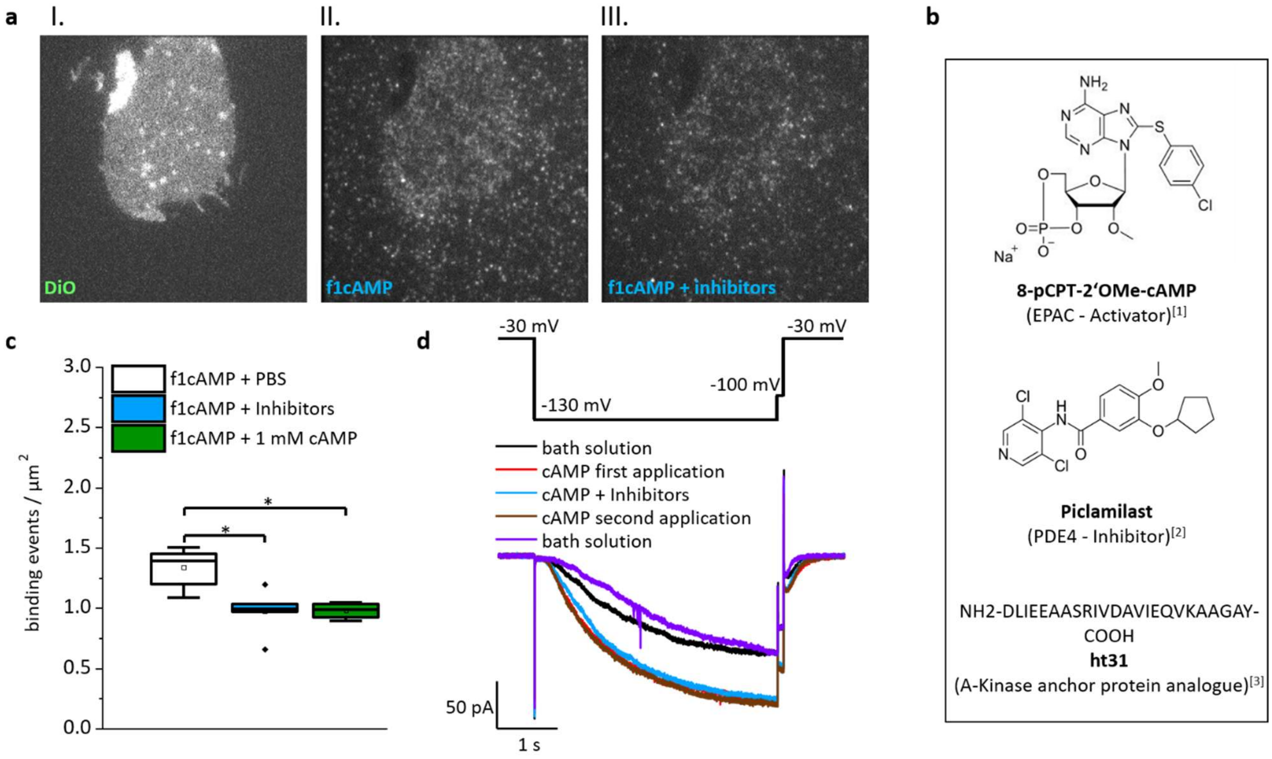


**Extended Data Fig. 5. Competitive inhibitors suppress unwanted binding of f1cAMP to endogenous cAMP binding membrane proteins. a,** Representative Micrographs. In step I, HEK293 cell membrane was stained with DiO which non-specifically stains membranes (488 nm excitation). In step II, binding of f1cAMP to membranes is observed (543 nm excitation). Signal also appears outside the membrane due to molecules randomly diffusing to the glass surface. Step III shows reduced binding of f1cAMP to the membrane while random diffusion to the glass surface is unaffected by additional competitive inhibitors. **b,** Structures of competitive inhibitors used to suppress binding to unwanted membrane proteins. **c,** Density of binding sites on HEK293 -derived membranes. The density of f1cAMP binding to the membrane is compared to f1cAMP binding in the presence of either the competitive Inhibitor mix or 1 mM cAMP as reference. * Indicates significant difference (ANOVA, Scheffé - test P<0.05). **d,** Patch-clamp recordings of mHCN2 channels showing different current amplitudes and kinetics depending whether cAMP was present or not. When competitive inhibitors were added no change in the currents was observed.

### Adaptation of the DISC – Algorithm

The DISC algorithm was taken from White and coworkers[14]. With our data, this algorithm often detected parts of white noise as additional steps. Therefore, the data were normalized on the total amplitude and DISC fits were accepted if all individual steps differ by at least 0.1 of the maximum amplitude from preceding levels. If this was not the case, the fit with n-1 steps was used instead and again checked if signals were at least 0.1 different from each other. The other DISC parameters were set to: alpha= 0.01, Viterbi=1, maximum number of states, k=5. We also identified a possible problem that could arise from using the algorithm on data with finite length. For example, if one channel binds one f1cAMP that would stay bound during the complete time of measurement, then we would have falsely identified the signal of that ligand as no ligand bound. Therefore, we would have identified additional ligands binding incorrectly as one, two and three ligands binding and by chance would never observe the fourth and final ligand. This would have drastically changed the outcome of our analysis.

Because of this risk, we implemented an additional quality control that compares the signal and noise of the lowest state in a trace to the background. Since the ligand signal was calculated out of a 3 × 3 area, we chose the ring of pixels surrounding that area with 2 pixels

distance as a surrogate for the background signal (Extended Data Fig. 6). Intensity of pixels contained in this ring was measured in each frame and the average and standard deviation were calculated. The Welch test was used to see if there are significant differences between the background level and the lowest identified state. If there were differences that were at least 95% of the average of binding levels of that trace, the lowest state would get declared as one ligand bound. The same was true if the difference was at least 95% of two times the average step size. Then the lowest state was declared to have two ligands bound. Analogously we treated the case of 3 bound ligands.
To ensure that this method does not generate artifacts, we simulated 10 traces per possible number of binding steps (40 traces in total) that were 10,000 frames long each. These idealized traces were combined with signal-to-noise ratios for each individual step from recorded data. In total, about 650 of these traces were then analyzed by the modified DISC algorithm. We compared the idealized traces to the original traces frame by frame. This resulted in about 85% of analyzed frames being correctly identified. However, upon investigation of individual traces it became clear, that either a trace was almost perfectly identified with a recovery rate, the fraction of points idealized back to its simulated state, above 95% or poorly matched the original because of the number of states in that trace was wrong. However, these traces were easily identifiable by eye. That is why for further analysis all traces were checked for plausibility prior to calculations by eye (see also Methods, Single molecule analysis).

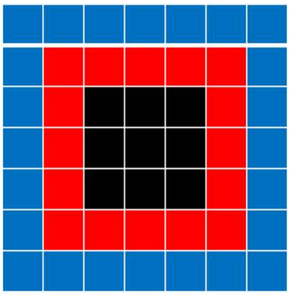

**Extended Data Fig. 6. Areas used for identification of signal and background**. The black area of pixels was used to calculate the ligand fluorescence signal, while the blue area was used to calculate the background signal. The red area was not used for calculations.
